# Exogenous Hormone Treatments Reveal Species-Specific Regulation of Individual Components of Root Architecture and Salt Ion Accumulation in Cultivated and Wild Tomatoes

**DOI:** 10.1101/2025.06.17.660158

**Authors:** Maryam Rahmati Ishka, Eric Craft, Miguel Pineros, Magdalena M. Julkowska

## Abstract

Hormonal signaling shapes plant architecture and salt stress responses, but its effects on root architecture and ion accumulation remain unclear. Here, we conducted a detailed analysis of how individual hormone treatments affect root architecture and ion accumulation under salt stress in tomato. The study focused on three tomato accessions with varying responses to salt stress. Our findings revealed distinct, species-specific hormonal effects. Auxin, ethylene, and gibberellin promoted lateral root development, yet their impacts on ion accumulation, particularly in Na⁺/K⁺ ratio, varied considerably. To explore the molecular basis of these differences, we examined Arabidopsis mutants for ethylene- and auxin-related genes, revealing novel components of hormone signaling involved in the salt stress response. Further, analysis of tomato mutants with impaired ethylene perception demonstrated that the *nr* mutant exhibits increased root growth and a higher shoot Na⁺/K⁺ ratio, largely due to reduced K⁺ retention. Our integrated physiological and genetic analysis reveals species-specific hormonal strategies can boost crop performance under salt stress.

## Introduction

Plant hormones are crucial in shaping plant architecture by regulating growth, development, and responses to environmental stimuli. Auxins are central to apical dominance and root initiation, while cytokinins promote cell division and shoot branching (Davies, 2010; Muraro *et al*., 2013; Rivas *et al*., 2022; Rowe *et al*., 2016; Casimiro *et al*., 2003; Marhavý *et al*., 2014). Gibberellins influence stem elongation and flowering, and abscisic acid (ABA) modulates stress responses and stomatal closure (Santner *et al*., 2009). Ethylene regulates leaf abscission and fruit ripening, and brassinosteroids are essential for cell expansion and vascular differentiation (Wang *et al*., 2025; Bleecker and Kende, 2000). These hormones function within intricate signaling networks, encompassing multiple logical circuits, allowing plants to adjust their structure to enhance resource utilization and survival. For example, the inhibition of primary root elongation by ABA depends on functional auxin transport and signaling pathways (X., Li *et al*., 2017). Yet, auxin-induced inhibition of root elongation occurs independently of ABA signaling (Thole *et al*., 2014). Insights into these hormonal pathways and dependencies offer opportunities to enhance crop productivity and stress tolerance through precise genetic or chemical strategies.

Recent studies have elucidated the role of plant hormones in regulating root growth and development under salt stress with species-specific insights. Abscisic acid (ABA) contributes to enhancing main root elongation and lateral root formation under saline conditions by modulating the expression of stress-responsive genes *AtABI4* and *AtABI5* in Arabidopsis (Shkolnik-Inbar *et al*., 2013). Cytokinins, on the other hand, suppress root growth (Werner *et al*., 2003), and in general high cytokinin levels are associated with decreased salt stress tolerance, potentially through interactions with ROS scavenging (Wang *et al*., 2015; Papon and Courdavault, 2022). In rice (*Oryza sativa*), auxin transporter *OsAUX1* was associated with increased lateral root development, enabling better nutrient acquisition under salt stress (Zhao *et al*., 2015). Medicago plants nodulated with the nitrogen-fixing bacterium *Sinorhizobium meliloti* RD64 exhibit higher levels of indole-3-acetic acid (IAA) in their nodules and roots compared to control plants, resulting in increased tolerance to salt stress (Bianco and Defez, 2009), suggesting exogenous application of IAA may enhance salt stress tolerance. GA3 treatments helped alleviate the effects of salt stress in maize roots, primarily by influencing ion accumulation and enhancing the activity of ROS-scavenging enzymes (Dinler, 2021). However, a detailed assessment of the specific components of root system architecture (RSA) is still missing. Ethylene generally promotes salinity tolerance in plants like Arabidopsis and maize (Riyazuddin *et al*., 2020). However, ethylene’s effects can depend on species of interest. For example, transgenic tobacco plants with suppressed ethylene production showed improved tolerance to salinity (Riyazuddin *et al*., 2020; Tavladoraki *et al*., 2012)), while an exogenous ethylene treatment in rice resulted in increased sensitivity to salinity stress (Yang *et al*., 2015). These contrasting findings underscore the need for systematic studies on how hormones impact individual components of salt tolerance. Dissecting plant RSA into three modular components: main root elongation, average lateral root growth, and increase in lateral root emergence (Julkowska *et al*., 2014) is a particularly well-suited system to systematically compare the effects of hormones on maintenance of root development, and can contribute to our understanding of plant stress resilience (Julkowska *et al*., 2017; Rahmati Ishka *et al*., 2024).

Although hormonal regulation of root systems under salt stress has been widely studied (Wang *et al*., 2015; Qin *et al*., 2019; Fu *et al*., 2019; Li *et al*., 2014; Shan *et al*., 2014; He *et al*., 2005; Moons *et al*., 1997; Lu *et al*., 2019), these studies were conducted across various plant species, developmental stages, and experimental setups, which limits the ability to directly apply their findings to other systems or contexts. Only a limited number of studies have investigated hormone application under salt stress beyond measuring primary root length (He *et al*., 2005; Lu *et al*., 2019), and even fewer have investigated its impact on ion accumulation (Fu *et al*., 2019; Li *et al*., 2014). Previously, we identified cultivated and wild tomato accessions with distinct RSAs under salt stress (Rahmati Ishka *et al*., 2024). The root-specific RNA-Seq suggested hormone signaling is an important component of salt stress responses that contributes to the maintenance of RSA (Rahmati Ishka *et al*., 2024). This observation leads us to the hypothesis that hormone treatments can be an effective strategy to boost the development of individual components of root architecture, regulate Na⁺ and K⁺ accumulation, and substantially contribute to overall salt stress tolerance. To test this hypothesis, we reexamined the root architecture of three tomato accessions with distinct salt tolerance strategies using agar plate-based assay. We found that although auxin, ethylene, and gibberellin can similarly impact RSA by increasing lateral root formation, their effects on ion accumulation and the resulting shoot Na⁺/K⁺ ratio vary significantly. To further investigate the underlying mechanisms, we analyzed Arabidopsis mutants for ethylene- and auxin-related genes identified in our root-specific RNA-Seq dataset (Rahmati Ishka et al., 2025), uncovering previously uncharacterized components of hormone signaling involved in salt stress responses. Additionally, experiments with tomato mutants defective in ethylene perception revealed that the *nr* mutant displays enhanced root growth and an elevated shoot Na⁺/K⁺ ratio, primarily due to impaired potassium retention. Together, these findings highlight the hormone-specific regulation of ion homeostasis under salt stress and provide a valuable framework for leveraging hormone pathways to improve crop resilience in saline environments—advancing sustainable agriculture and global food security.

## Methods

### Agar-based plate experiments

Tomato seeds were sterilized for 10 min with 50% bleach and rinsed five times using autoclaved milli-Q water, and germinated on ¼ strength Murashige and Skoog (MS) medium containing 0.5% (w/v) sucrose, 0.1% (w/v) 4-morpholineethanesulfonic acid (MES), and 1% (w/v) agar, with pH adjusted to 5.8 with KOH. After 24 h of vernalization at 4^◦^C in the dark, the plates were placed in the Conviron growth chamber with the light intensity of 130–150 µmol x m^-2^ x s ^-1^ in a 20 h light / 4 h dark cycle at 25 ^◦^C day / 20 ^◦^C night and 60% humidity. 4 days after germination, the seedlings were transferred to 1/4 MS media with and without 100 mM NaCl or other treatments, as indicated in the figures. Each plate contained one tomato seedling for RSA analysis. The plates were scanned using EPSON scanner for 5 consecutive days, starting from day 5 after germination.

Arabidopsis seeds were processed as described in (Ishka *et al*., 2024). Briefly, the seeds were sterilized for 10 min with 50% bleach and, rinsed five times using milli-Q water, and germinated on ½ strength Murashige and Skoog (MS) medium, with PH adjusted to 5.8 with KOH. After 24 h of vernalization at 4^◦^C in the dark, the plates were placed in the Conviron growth chamber with the light intensity of 130–150 µmol x m^−2^ x s ^−1^ in a 16 h light / 8 h dark cycle at 21^◦^C and 60% humidity. 4 days after germination, the seedlings were transferred to ½ MS media supplemented with 0 or 75 mM NaCl as indicated. The plates were scanned using EPSON scanner every other day, starting from 4 days after germination until the plants were 14 days old.

To analyze RSA traits from the scanned plate images, we used SmartRoot plugin (Lobet *et al*., 2011) in ImageJ to manually trace the root and extract root-related features in the CSV format, followed by data analysis in R.

For analysis of RSAs that included hormonal treatments, we used the following concentrations: For ethylene, we used ethylene precursor, 1-aminocyclopropane-1-carboxylic acid (ACC), in the ¼ MS plates as described above, supplemented with 1 or 5 µM ACC. For IAA, the MS media was supplemented with 0.05 or 1 µM IAA. GA3 stock was made in absolute ethanol and was used in 1 or 5 µM final concentration in the MS media. ABA stock was prepared in absolute ethanol, and 1 or 10 uM was used as the final concentration. Trans-zeatin was used for cytokinin treatment. The stock was prepared in DMSO and used in the final concentration of 1 or 10 µg/ml. All hormones were added after filter-sterilization into the autoclaved cool media.

### Soil experiments

Unless otherwise stated, the tomato seeds were germinated in 1/4 MS media, as described for the agar-based plate assay above. On day 5, seedlings were transferred to 1/4 MS plates containing either 0 or 100 mM NaCl for one week. Subsequently, the seedlings were transplanted into Cornell OSMOCOT soil containing 0 or 100 mM NaCl with 50% water holding capacity (WHC) and placed in the walk-in growth chamber with the 16 h light / 8 h dark period, 25 ^◦^C day / 20 ^◦^C night throughout the growth period and 60% humidity. The water holding capacity was maintained at 50% throughout the experiment by manually measuring the pot weight daily and adjusting it with the water. Using the PhenoCage setup (Yu *et al*., 2024), plant images were captured every other day, starting from 28 days after stress and continuing for 8 days. Daily growth rates were calculated for each plant based on the estimated shoot growth area obtained from the 7-side view images.

For those soil experiments that included foliar hormonal treatments in tomato, we used the following concentrations: For ACC and IAA foliar sprays, we used 1 µM for both hormones. Water was used as a mock treatment for ACC in the soil experiment. Because IAA stock was dissolved in KOH solution, the same amount of KOH was used in the mock treatment. Spraying was done manually using spry bottles on leaves, starting from the day of transplanting into the soil for 5 consecutive days, followed by twice per week for a total of 4 weeks for tomato, and until maturation and seed formation for Arabidopsis plants.

### Shoot size and evapotranspiration measurements

We used Side-view imaging using PhenoCage as described here (Yu *et al*., 2024)to measure the shoot growth in soil. The collected images were processed using PlantCV pipeline (Gehan *et al*., 2017), followed by the data analysis in R.

The plant evapotranspiration was measured manually by measuring the pot’s weight every day and watering them to their target weight.

### Electrolyte leakage analysis

The electrolyte leakage measurement was done as described in (Ishka *et al*., 2024). In brief, for each plant, three leaf discs were incubated in 2 ml of distilled water for 24 h at the light at room temperature with gentle shaking, followed by measuring the initial electrolyte leakage using a conductivity meter (Horiba LAQUAtwin EC-11 Compact Conductivity Meter, # 3999960125). The samples were then subjected to 80 ^◦^C for 2 h to release the total electrolyte and cooled at room temperature for 24 h. Final electrolyte leakage was measured the following day. The electrolyte leakage percentage was calculated by dividing initial conductivity by final conductivity * 100, followed by data analysis in R.

### Elemental analysis

The elemental analysis was performed using Inductively Coupled Plasma Atomic Emission Spectroscopy (ICP-AES). For analysis of Na^+^ and K^+^ ions in roots and shoots of plate-grown seedlings, the root and shoot tissues were harvested after 10 days of treatments, as indicated in the figures. For soil-grown plants, the samples were harvested at the end of week four. All samples were rinsed in milli-Q water and collected into paper bags and dried at 60 ^◦^C oven for one week. Subsequently, the dry weight was recorded. Samples were digested in double distilled HNO3, followed by the addition of 60/40 nitric/perchloric acid, continuing incubation at 150 °C. Samples were processed for ICP-AES analysis using a Thermo iCap 7000 ICP-AES after being diluted to 10 ml with deionized water. Ion content was calculated per dry weight for each sample, followed by data analysis in R.

### Screening tomato and Arabidopsis hormone-related mutants

Two ethylene-related tomato mutants, provided by Professor Jim Giovannoni’s lab at the Boyce Thompson Institute (BTI), were examined in this study. The *never-ripe* (*nr*) mutant, in the Ailsa Craig (AC) background, carries a dominant mutation in an ethylene receptor, resulting in ethylene insensitivity, delayed fruit ripening, and prolonged green fruit retention (Lanahan et al., 1994). The *non-ripening ethylene insensitive* (*nei*) mutant, in the M82 background, has a recessive mutation in the EIN2 ortholog, causing ethylene insensitivity (Guo et al., 2018).

The following T-DNA insertion alleles were used for ethylene-related mutants: for *EIL4* (AT5G10120), *eil4-1* (SAIL_623_H07, Basta resistant) and *eil4-2* (SALKseq_034500.1, kanamycin resistant); for EGY2 (AT5G05740), egy2-1 (SAILseq_735_G12.2, Basta resistant) and egy2-2 (SALK_001991, kanamycin resistant) were used. The following T-DNA insertion alleles were used for auxin-related mutants: for *PILS5* (AT2G17500), *pils5-1* (SALKseq_121078.1, kanamycin resistant) and *pils5-3* (SALK_072996C, kanamycin resistant); for SAUR45 (AT2G36210), saur45-1 (SAIL_274_H04, basta resistant) and saur45-2 (SALK_041272C, kanamycin resistant) were used. All T-DNA insertion alleles were in the background of Col-0 accession. The primers used for genotyping PCR are shown in **Table S1**.

## Results

### Cultivated and wild-tolerant tomatoes prioritize different aspects of lateral root development under salt stress

Based on RSA responses to salt stress observed in a wild tomato diversity panel (Rahmati Ishka et al., 2025), we selected three tomato accessions for detailed analysis: LA2540, a wild salt-tolerant accession (hereafter referred to as wild tolerant); LA1371, a wild salt-sensitive accession (wild sensitive); and LA1511, a *Solanum lycopersicum* accession (cultivated tomato). Under salt stress, main root elongation was inhibited only in the wild-sensitive accession, while the main root of the wild tolerant and cultivated accessions remained unaffected (**Fig. 1A**). In terms of lateral root length, a significant reduction was observed only in the cultivated tomato, whereas both wild accessions maintained unchanged lateral root lengths under salt stress (**Fig. 1B**). For lateral root number (**Fig. 1C**), the cultivated tomato showed no difference with or without salt, but both wild accessions exhibited a significant reduction under salt stress. These findings are consistent with previously reported phenotypes, where cultivated tomato tends to prioritize lateral root emergence, whereas wild salt-tolerant tomato emphasizes lateral root elongation under salt stress—highlighting distinct adaptive strategies among different accessions in response to salinity.

**Figure 1.**
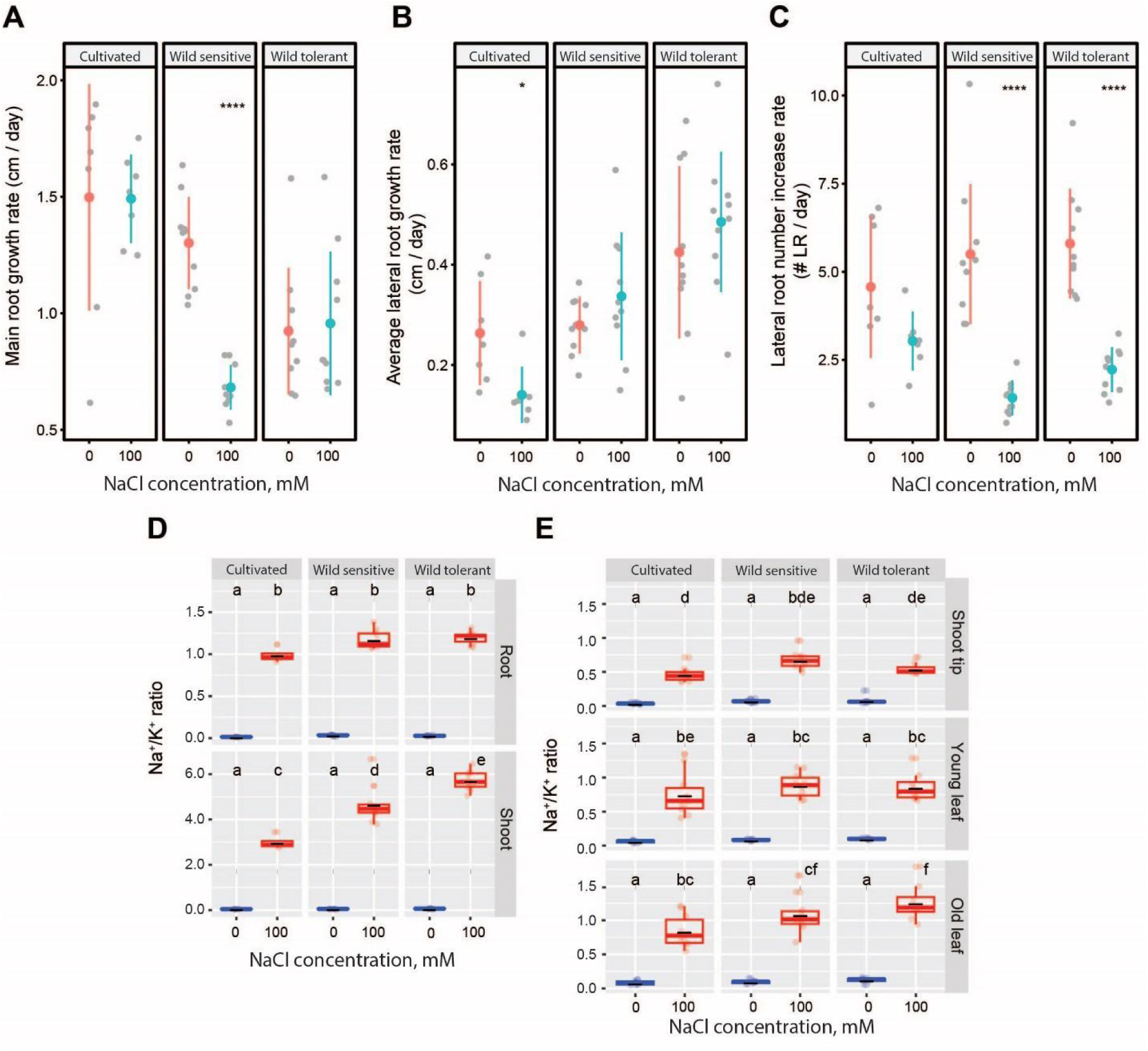
Wild tolerant and cultivated tomatoes have different salt tolerant features regarding lateral root development under the salt. Root system architecture analysis of three tomato accessions under 0 or 100 mM concentrations of NaCl are shown for **(A)** main root length, **(B)** lateral root growth rate, and lateral root numbers. **(D)** Na^+^/K^+^ ratio in root and shoot of different accessions after 10 days on treatment plates. **(E)** Na⁺/K⁺ ratio in three distinct shoot tissues of four-week-old plants after two weeks of salt stress in soil. Lines in (A-C) graphs represent median values. The asterisks above the graphs in (A-C) indicate significant differences between control and salt stress conditions, as determined by the Student‘s t-test: *P < 0.05, **P < 0.01, ***P < 0.001, and ****P < 0.0001. (D-E) Statistical analysis was done by comparison of the means for all pairs using Tukey HSD test. Levels not connected by the same letter are significantly different (P < 0.05).

We performed ICP-AES analysis after 10 days of salt exposure (Fig. 1D, Fig. S1A-B) to assess ion accumulation. Salt stress exposure resulted in increased and decreased Na^+^ and K^+^ accumulation, respectively, with no significant differences between the accessions (**Fig. S1A-B**). Although all accessions maintained a similar Na^+^/K^+^ ratio in the roots, cultivated tomatoes exhibited a significantly lower Na^+^/K^+^ ratio in the shoot compared to both wild accessions. This indicates that cultivated tomatoes may be more efficient at restricting Na⁺ translocation to the shoot, whereas wild accessions likely tolerate higher shoot Na⁺ levels, reflecting distinct tissue-level strategies for coping with salt stress.

To further explore the responses of these accessions to salt stress beyond the seedling stage, we evaluated their performance in soil (**Fig. S1C-H**). Salt stress caused a significant reduction in shoot size (**Fig. S1C**) and shoot fresh weight (**Fig. S1D**) across all accessions tested. We also analyzed plant evapotranspiration and found no significant differences among the accessions under control conditions (**Fig. S1E**). However, salt stress led to a significant increase in evapotranspiration in all accessions compared to the control (**Fig. S1E**). Interestingly, while the cultivated and wild-tolerant tomatoes showed similar evapotranspiration rates under salt stress, the wild-sensitive tomato exhibited a significantly higher rate, highlighting its greater sensitivity to salt stress (**Fig. S1E**).

To assess cellular integrity during salt stress, we measured changes in cell membrane integrity by evaluating ion leakage in leaf discs at the end of the experiment (**Fig. S1F**). Salt stress significantly increased ion leakage across all accessions compared to the control in both old and young leaves. In old leaves, the wild-tolerant tomato exhibited a significant increase in ion leakage compared to the cultivated tomato, but the increase was not significant compared to the wild-sensitive tomato (**Fig. S1F**). Similarly, in young leaves, the wild-tolerant tomato showed a non-significant increase in ion leakage compared to both the cultivated and wild-sensitive tomatoes (**Fig. S1F**).

We further analyzed shoot ion accumulation in the selected accessions using ICP-AES after two weeks of salt stress exposure (**Fig. 1E, Fig. S1G-H**). Salt stress caused a gradual increase in Na⁺ from the shoot tip to the old leaf in all accessions, with the wild-tolerant type showing the highest Na⁺ in old leaves (**Fig. S1G**). K⁺ levels stayed mostly stable, with only slight, non-significant changes across tissues and accessions (**Fig. S1H**). The Na⁺/K⁺ ratio also increased along the shoot and was highest in the old leaves of the wild-tolerant accession (**Fig. 1E**). Together, these findings are consistent with the agar-based experiment, where the wild-tolerant accession also showed a higher Na⁺/K⁺ ratio in shoot (**Fig. 1D**), further supporting the idea that it may compartmentalize excess Na⁺ in older leaves to protect younger and developing tissues during salt stress.

### Auxin treatment reduced root length, increased lateral root numbers, and elevated shoot Na^+^/ K^+^ ratio in wild tomatoes but not cultivated tomato

To examine how auxin treatment influences the RSA of the three tomato accessions with distinct root architecture, we treated plants with indole-3-acetic acid (IAA), as it is the naturally occurring auxin in plants, with and without salt stress (**Fig. 2, Fig. S2**). IAA treatment significantly reduced both main root length (**Fig. 2A-B**) and lateral root length (**Fig. 2C**), while simultaneously increasing lateral root numbers (**Fig. 2D**) across all accessions and conditions. These results suggest that IAA promotes lateral root initiation at the expense of root elongation, regardless of salt stress or genetic background.

**Figure 2.**
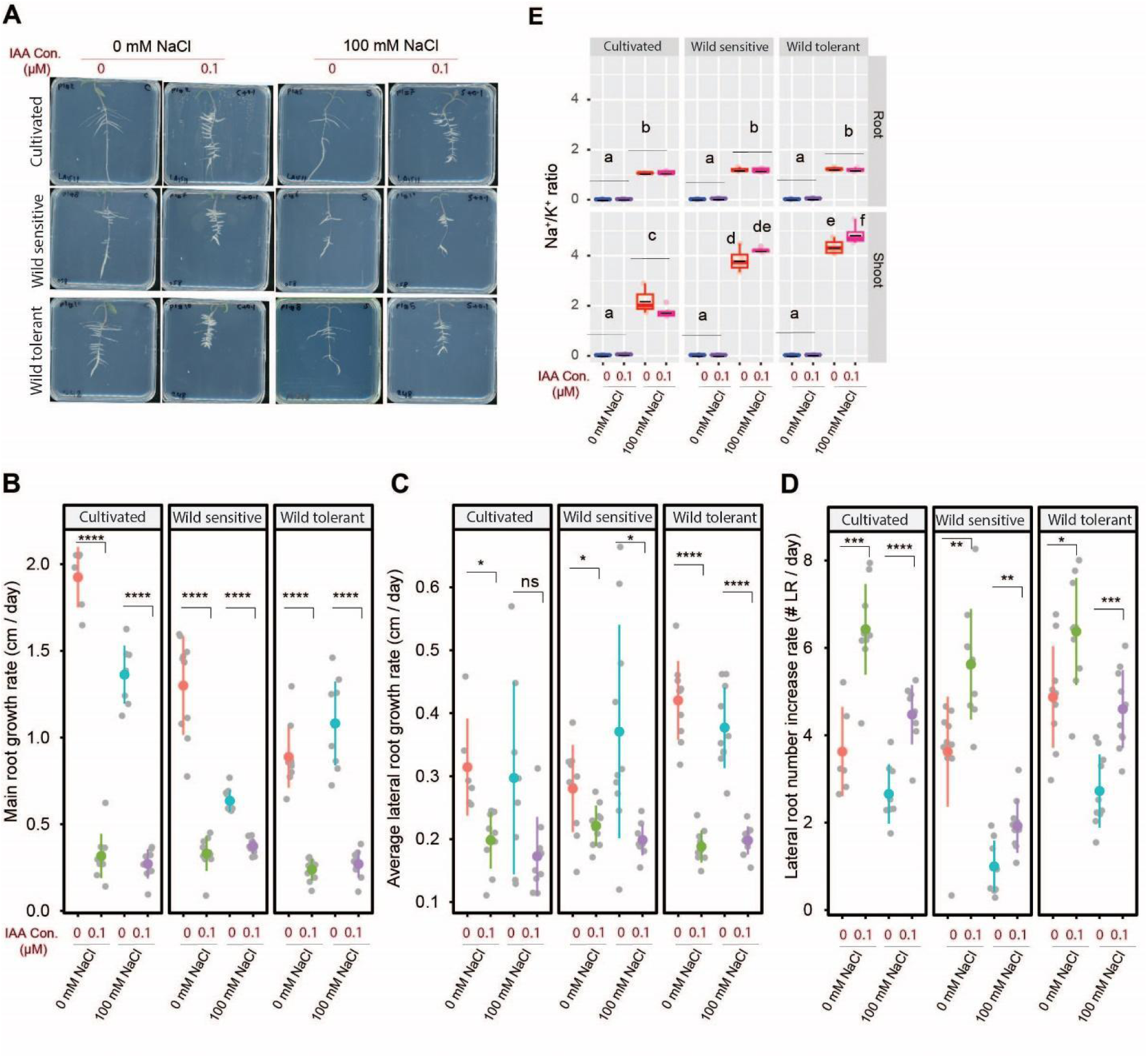
IAA treatment reduces root growth while increasing lateral root number across all accessions and conditions but only raises the Na⁺/K⁺ ratio in shoots of wild-tolerant tomatoes. **(A)** Shown are representative images of 9-day-old tomato seedlings that experienced 5 days of treatment with 0 or 100 mM concentrations of NaCl supplemented with or without various concentrations of IAA, as indicated in the figure. Root system architecture analysis is shown here for the main root growth rate **(B)**, average lateral root growth **(C)**, and number of lateral roots **(D)**. The asterisks above the graphs in (B-D) indicate significant differences between 0 and 0.1 µM IAA under either control or salt stress conditions by the Student‘s t-test: *P < 0.05, **P < 0.01, ***P < 0.001, and ****P < 0.0001. Na^+^/K^+^ ratio **(E)** of root and shoot of different accessions after 10 days on treatment plates. Each dot in (E) represents individual replicate per accession. Lines in (E) represent mean values. Statistical analysis was done by comparison of the means for all pairs using Tukey–Kramer HSD test for (E). Levels not connected by the same letter are significantly different (P < 0.05).

To assess ion accumulation under IAA treatment, we measured Na⁺, K⁺, and their ratio in roots and shoots after 10 days of salt exposure (**Fig. 2E, Fig. S2A-B**). Salt stress increased Na⁺ and decreased K⁺ in all accessions, with no major differences (**Fig. S2A-B**). IAA caused a slight, non-significant rise in root Na⁺ across accessions but significantly increased shoot Na⁺ only in the wild-tolerant tomato (**Fig. S2A**). For K⁺, IAA reduced root K⁺ under control conditions only in the wild-tolerant type, with no effect under salt (**Fig. S2B**). In shoots, IAA lowered K⁺ in the cultivated tomato under control but had no effect on wild accessions or under salt stress (**Fig. S2B**). IAA treatment did not affect the Na⁺/K⁺ ratio in roots across accessions, but it significantly increased the ratio in the shoots of the wild-tolerant tomato, with a slight, non-significant increase in the wild-sensitive type and no change in the cultivated tomato (**Fig. 2E**). These results suggest that IAA influences shoot ion homeostasis in an accession- and tissue-specific manner, particularly enhancing Na⁺ accumulation in wild tomatoes under salt stress.

### ACC treatment enhanced lateral root formation under salt stress in wild tomatoes, without altering Na^+^/ K^+^ ratio

To assess the effect of ethylene on root architecture, we treated the three tomato accessions with ACC, an ethylene precursor, with and without salt stress (**Fig. 3, Fig. S3**). ACC significantly reduced main root length in all accessions under all conditions (**Fig. 3A-B**). For lateral root length, only the higher ACC concentration reduced it under control, while both concentrations inhibited it under salt in wild accessions; in cultivated tomato, only the lower ACC was inhibitory (**Fig. 3C**). Lateral root number responses were accession-specific. Under control, lower ACC reduced it in cultivated and wild-sensitive tomatoes, while wild-tolerant remained unaffected. Under salt, lower ACC reduced lateral roots in cultivated but increased them in both wild accessions, with wild-tolerant showing the strongest response at high ACC (**Fig. 3D**). These results suggest ACC modulates root architecture in a concentration- and genotype-specific manner, particularly enhancing lateral root formation under salt stress in wild tomatoes.

**Figure 3.**
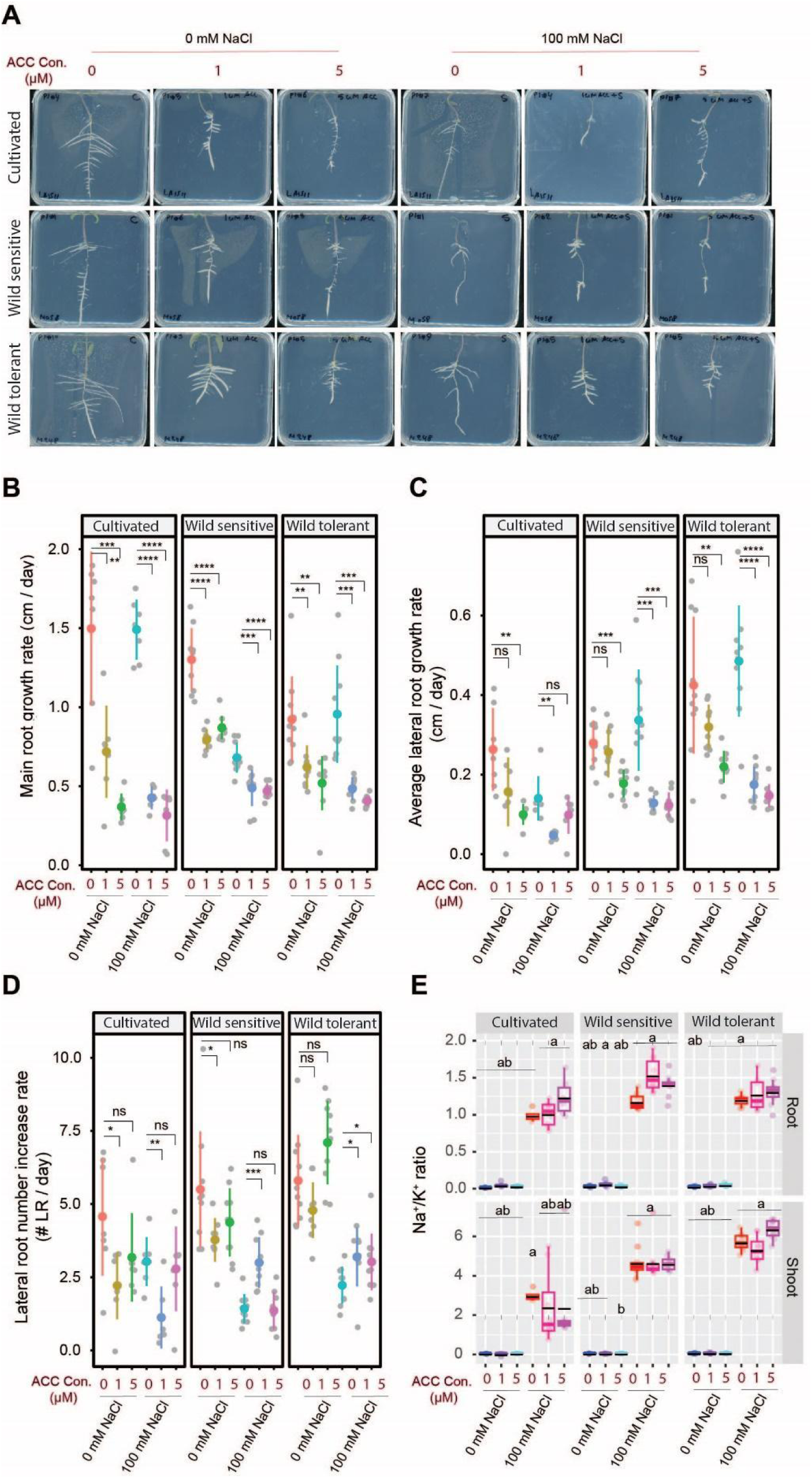
ACC treatment significantly reduces main root and lateral root length while increasing lateral root numbers without altering Na^+^/K^+^ ratio. **(A)** Shown are representative images of 9-day-old tomato seedlings that experienced 5 days of treatment with 0 or 100 mM concentrations of NaCl supplemented with or without various concentrations of ACC as indicated in the figure. Root system architecture analysis are shown here for main root growth rate **(B)**, average lateral root growth **(C)**, and number of lateral roots **(D)**. The asterisks above the graphs in (B-D) indicate significant differences between ACC and no ACC treatments under either control or salt stress conditions by the Student‘s t-test: *P < 0.05, **P < 0.01, ***P < 0.001, and ****P < 0.0001. Na^+^/K^+^ ratio **(E)** of root and shoot of different accessions after 10 days on treatment plates. Each dot in (E) represents individual replicate per accession. Lines in (E) represent mean values. Statistical analysis was done by comparison of the means for all pairs using Tukey–Kramer HSD test for (E). Levels not connected by the same letter are significantly different (P < 0.05).

To assess ion accumulation under ACC treatment, we measured Na⁺, K⁺, and their ratio in roots and shoots after 10 days of salt exposure (**Fig. 3E, Fig. S3A-B**). Salt increased Na⁺ and reduced K⁺ across accessions, with no major differences across accessions. ACC slightly reduced Na⁺ in most cases, except in the shoot of wild-sensitive tomato, where Na⁺ levels remained unchanged (**Fig. S3A**). K⁺ levels were unaffected by ACC under all conditions (**Fig. S3B**). Overall, ACC had no detectable impact on the Na⁺/K⁺ ratio in roots or shoots (**Fig. 3E**). These results suggest that ACC treatment has a limited impact on shoot and root Na⁺ and K⁺ homeostasis under salt stress, at least under the conditions tested.

### ABA treatment inhibited root system architecture and increased shoot Na^+^/ K^+^ ratio, except in wild-tolerant tomato

Among the hormones evaluated for their impact on root system architecture and salt stress resilience, abscisic acid (ABA), a key stress-responsive hormone, was included. Under control conditions, ABA treatment significantly reduced main root length in both cultivated and wild-tolerant tomatoes, while the wild-sensitive accession remained unaffected (**Fig. 4A-B**). Under salt stress, ABA treatment led to a significant reduction in main root length across all accessions, except in the wild-sensitive tomato, where a low concentration of ABA notably promoted main root growth (**Fig. 4B**). Across all accessions and treatment conditions, ABA consistently suppressed lateral root length (**Fig. 4C**). Furthermore, ABA reduced lateral root number under control conditions in all tomato accessions (**Fig. 4D**). Under salt stress reduction in lateral root number was observed only in the cultivated tomato, with no significant effect in the two wild accessions (**Fig. 4D**). These findings suggest that ABA plays a predominantly inhibitory role in shaping root architecture, with distinct accession-specific responses under salt stress.

**Figure 4.**
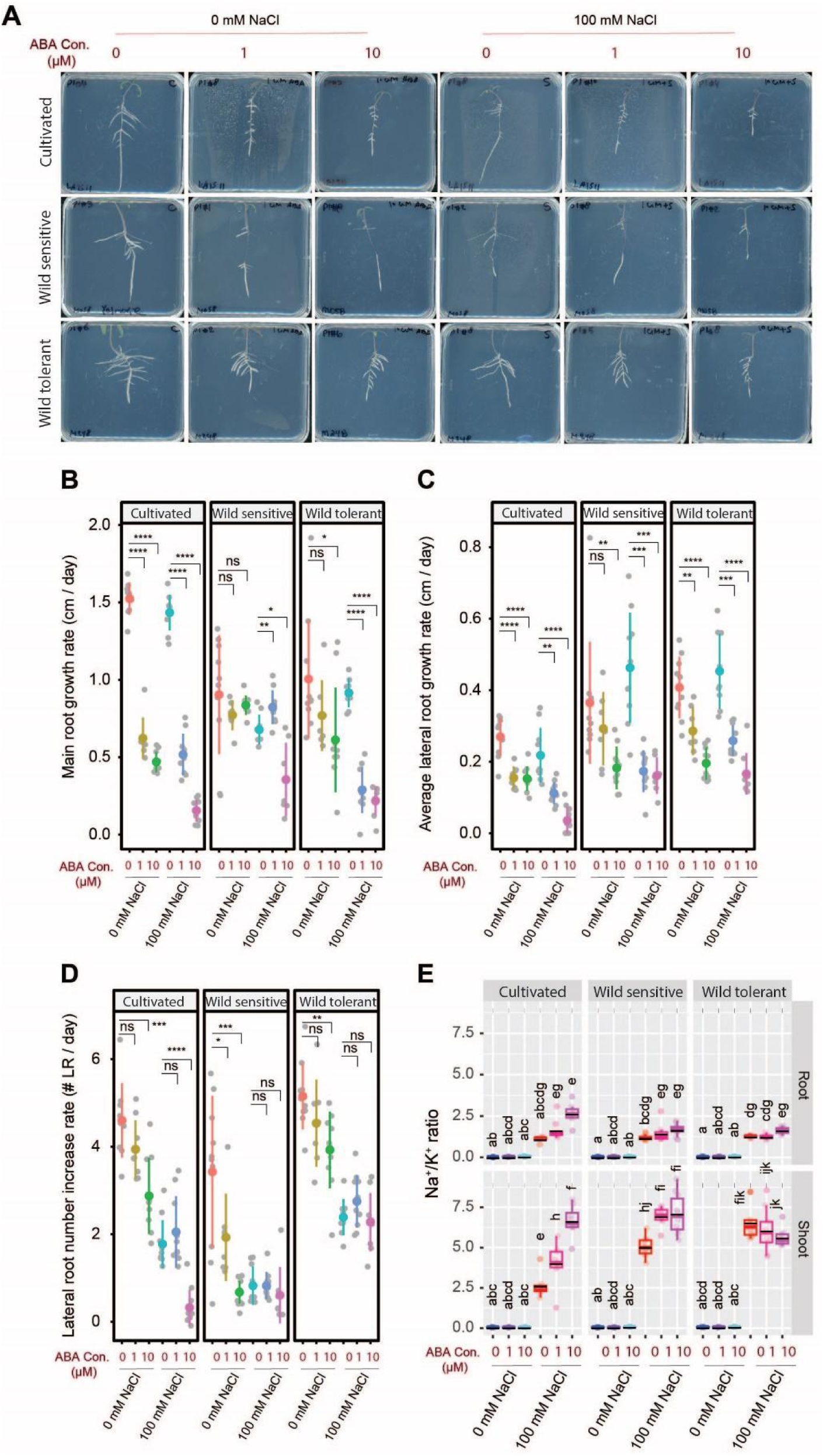
ABA treatment inhibits root growth, while increasing Na/K ratio in all accessions but not wild tolerant. **(A)** Shown are representative images of 9-day-old tomato seedlings that experienced 5 days of treatment with 0 or 100 mM concentrations of NaCl supplemented with or without various concentrations of ABA as indicated in the figure. Root system architecture analysis are shown here for main root growth rate **(B)**, average lateral root growth **(C)**, and number of lateral roots **(D)**. The asterisks above the graphs in (B-D) indicate significant differences between ABA and no ABA treatments under either control or salt stress conditions by the Student‘s t-test: *P < 0.05, **P < 0.01, ***P < 0.001, and ****P < 0.0001. Na^+^/K^+^ ratio **(E)** of root and shoot of different accessions after 10 days on treatment plates. Each dot in (E) represents individual replicate per accession. Lines in (E) represent mean values. Statistical analysis was done by comparison of the means for all pairs using Tukey–Kramer HSD test for (E). Levels not connected by the same letter are significantly different (P < 0.05).

To evaluate the impact of ABA treatment on ion accumulation, we analyzed Na^+^, K^+^, and their ratio in the roots and shoots of seedlings (**Fig. 4E, FigS4A-B**). Our findings showed that different ABA concentrations had varying effects on Na^+^ accumulation, depending on the accession. While higher ABA concentrations did not affect root Na^+^ content in cultivated and wild-tolerant tomatoes, they seem to reduce Na^+^ accumulation in the shoot compared to salt stress alone, although non-significantly (**Fig. S4A**). ABA treatment decreased K^+^ content in all accessions in both roots and shoots under control conditions (**Fig. S4B**). Under salt stress, ABA reduced K^+^ content in both roots and shoots of cultivated tomato but had no effect on wild tomatoes (**Fig. S4B**). ABA treatment led to a non-significant increase in the Na⁺/K⁺ ratio in roots and a significant increase in shoots of cultivated tomato (**Fig. 4E**). The wild-sensitive accession maintained the same Na^+^/K^+^ ratio in roots under ABA treatment but exhibited a significantly higher ratio in shoots compared to salt stress alone (**Fig. 4E**). In contrast, the wild-tolerant accession maintained a nearly unchanged Na^+^/K^+^ ratio in both roots and shoots under ABA treatment relative to salt stress alone (**Fig. 4E**). These results indicate that ABA modulates ion homeostasis under salt stress in an accession- and tissue-specific manner, primarily by limiting K⁺ retention in cultivated tomatoes, while this effect is absent in wild accessions.

### Gibberellin increased lateral root numbers but reduced the shoot Na⁺/K⁺ ratio under salt stress

Because gibberellin (GA) is known to affect root development under salt stress by interfering with auxin signaling (Lv *et al*., 2018), we decided to test its impact on RSA and salt stress resilience in tomato varieties here. We used GA₃ as it is one of the most commonly applied synthetic gibberellins. Under control conditions, GA₃ reduced the main root length in all accessions, except for a low concentration that significantly increased it in cultivated tomato (**Fig. 5A-B**). Under salt stress, GA₃ promoted main root growth across all accessions, though high GA₃ significantly reduced root length in cultivated tomato (**Fig. 5B**). GA₃ decreased lateral root length in cultivated tomato under both conditions but had contrasting effects on wild accessions—no effect under control, yet stimulation under salt stress, with high GA₃ most effective in wild sensitive and low GA₃ in wild tolerant tomatoes (**Fig. 5C**). Lateral root number, remained mostly unchanged under control, except for a decrease in wild tolerant tomato with low GA₃ (**Fig. 5D**). Under salt stress, GA₃ increased lateral root numbers in all accessions, with the most effective concentration varying by accession (**Fig. 5D**). These findings suggest that GA₃ modulates root architecture in a concentration- and accession-dependent manner, with enhanced promotive effects under salt stress.

**Figure 5.**
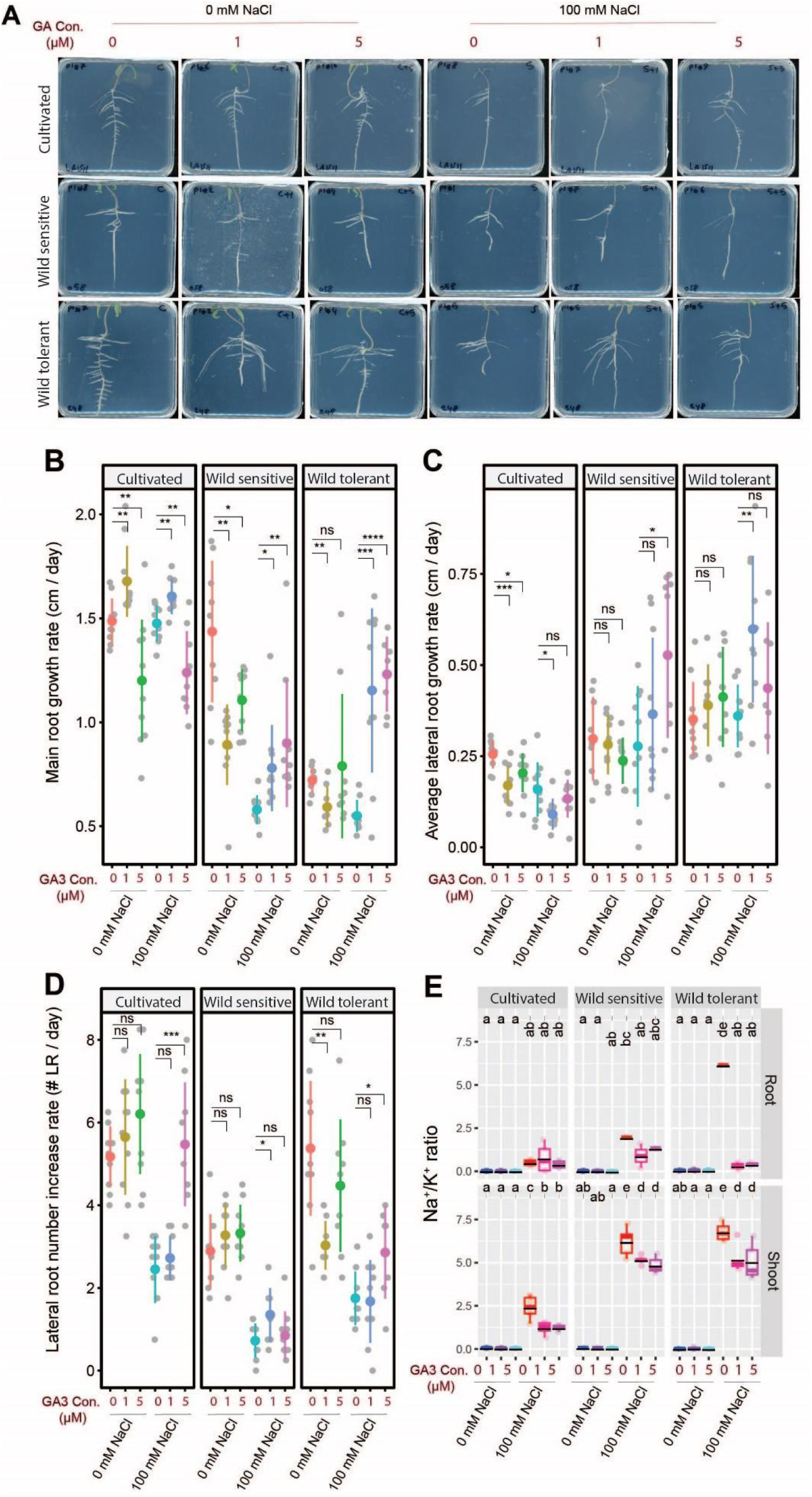
GA3 treatment increases lateral root numbers while decreasing shoot Na/K ratio in all accessions. **(A)** Shown are representative images of 9-day-old tomato seedlings that experienced 5 days of treatment with 0 or 100 mM concentrations of NaCl supplemented with or without various concentrations of GA3, as indicated in the figure. Root system architecture analysis is shown here for main root growth rate **(B)**, average lateral root growth **(C)**, and number of lateral roots **(D)**. The asterisks above the graphs in (B-D) indicate significant differences between GA3 and no GA3 treatments under either control or salt stress conditions by the Student‘s t-test: *P < 0.05, **P < 0.01, ***P < 0.001, and ****P < 0.0001. Na^+^/K^+^ ratio **(E)** of root and shoot of different accessions after 10 days on treatment plates. Each dot in (E) represents individual replicate per accession. Lines in (E) represent mean values. Statistical analysis was done by comparison of the means for all pairs using Tukey–Kramer HSD test for (E). Levels not connected by the same letter are significantly different (P < 0.05).

To evaluate the impact of GA₃ treatment on ion accumulation, we analyzed Na^+^, K^+^, and their ratio in the roots and shoots of seedlings (**Fig. 5, Fig. S5A-B**). While considerable variation in Na⁺ and K^+^ accumulations was observed across all conditions, GA₃ treatment reduced root Na⁺ levels in all accessions at one or both concentrations (**Fig. S5A**) and enhanced K⁺ retention in roots and shoots under salt stress (**Fig. S5B**). While cultivated tomato maintained a similar root Na⁺/K⁺ ratio (**Fig. 5E**), wild-sensitive and wild-tolerant accessions showed a non-significant and significant reduction, respectively (**Fig. 5E**). GA₃ significantly lowered the shoot Na⁺/K⁺ ratio in all accessions (**Fig. 5E**). These results suggest that GA₃ promotes salt tolerance by enhancing Na⁺ exclusion and K⁺ retention in an accession- and tissue-specific manner.

### Cytokinin treatment suppressed all aspects of root system architecture across all accessions while elevating the shoot Na**⁺**/K**⁺** ratio specifically in cultivated tomato

To assess the effect of cytokinin on root architecture and salt resilience, we treated the three tomato accessions with trans-zeatin, a biologically active form of cytokinin, with and without salt stress (**Fig. 6, Fig. S6**). Our results showed that cytokinin treatment significantly reduced main root length in all accessions under both control and salt conditions, except for the low concentration, which had no effect on wild tomatoes (**Fig. 6A-B**). Similarly, cytokinin significantly decreased both lateral root length and number across all accessions and treatments, except for the wild-sensitive tomato under salt stress, which remained unaffected compared to salt treatment alone (**Fig. 6C-D**). These findings suggest that cytokinin generally suppresses root growth under both normal and saline conditions.

**Figure 6.**
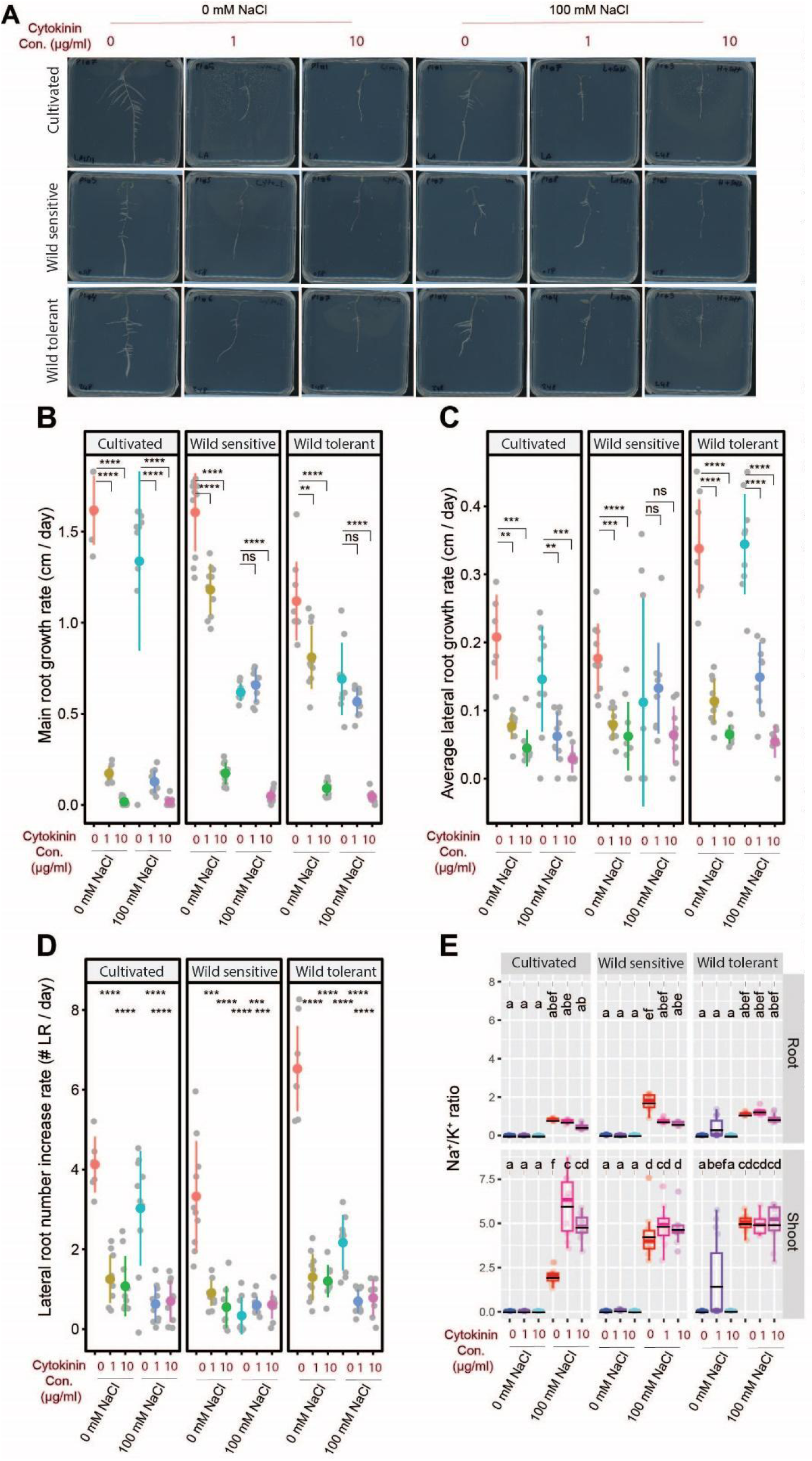
Cytokinin treatment inhibits root growth while increasing shoot Na/K ratio in cultivated tomato. **(A)** Shown are representative images of 9-day-old tomato seedlings that experienced 5 days of treatment with 0 or 100 mM concentrations of NaCl supplemented with or without various concentrations of GA3 as indicated in the figure. Root system architecture analysis is shown here for the main root growth rate **(B)**, average lateral root growth **(C)**, and number of lateral roots **(D)**. The asterisks above the graphs in (B-D) indicate significant differences between cytokinin and no cytokinin treatments under either control or salt stress conditions by the Student‘s t-test: *P < 0.05, **P < 0.01, ***P < 0.001, and ****P < 0.0001. Na^+^/K^+^ ratio **(E)** of root and shoot of different accessions after 10 days on treatment plates. Each dot in (E) represents individual replicate per accession. Lines in (E) represent mean values. Statistical analysis was done by comparison of the means for all pairs using Tukey–Kramer HSD test for (E). Levels not connected by the same letter are significantly different (P < 0.05).

To evaluate the impact of trans-zeatin treatment on ion accumulation, we analyzed Na^+^, K^+^, and their ratio in the roots and shoots of seedlings (**Fig. 6, Fig. S6A-B**). Analysis of Na⁺ content revealed accession-specific responses to cytokinin treatment (**Fig. 6, Fig. S6A**). Cytokinin had no effect on root Na⁺ accumulation but significantly increased shoot Na⁺ levels in cultivated and wild-sensitive tomatoes, with no impact on the wild-tolerant accession (**Fig. S6A**). Root K⁺ content remained unchanged under all conditions, while shoot K⁺ levels decreased under control conditions in all accessions but were unaffected under salt stress (**Fig. S6B**). Cytokinin reduced the root Na⁺/K⁺ ratio only in the wild-sensitive accession, while it significantly and non-significantly increased the shoot Na⁺/K⁺ ratio in cultivated and wild-sensitive tomatoes, respectively, with no effect in wild-tolerant plants (**Fig. 6E**). These results indicate that cytokinin disrupts shoot ion homeostasis under salt stress in an accession-dependent manner, mainly by promoting Na⁺ accumulation.

### IAA raises shoot Na⁺/K⁺ ratios in both tomato types, while ACC lowers it only in wild-tolerant tomatoes under early salt stress

Based on early seedling establishment assays in plates, we observed that IAA significantly increased the Na⁺/K⁺ ratio, while ACC treatment had no impact on ionic balance under salt stress. To investigate how these hormones affect salt stress responses beyond the plate-based seedling stage, we selected IAA and ACC for further study by spraying them onto the leaves of soil-grown plants from two salt-tolerant tomato accessions, cultivated and wild tolerant, representing distinct salt tolerance strategies. By examining their effects during later developmental stages, we aimed to uncover whether early ionic trends persist and how these hormones differentially regulate growth and ion homeostasis over time.

The foliar application of IAA significantly reduced shoot size under control conditions in both accessions compared to the mock treatment (KOH solution without IAA). However, under salt stress conditions, shoot size was not affected by IAA application (**Fig. 7A, Fig. S7A**). These results suggest that the growth-inhibitory effect of IAA is specific to non-stress conditions.

**Figure 7.**
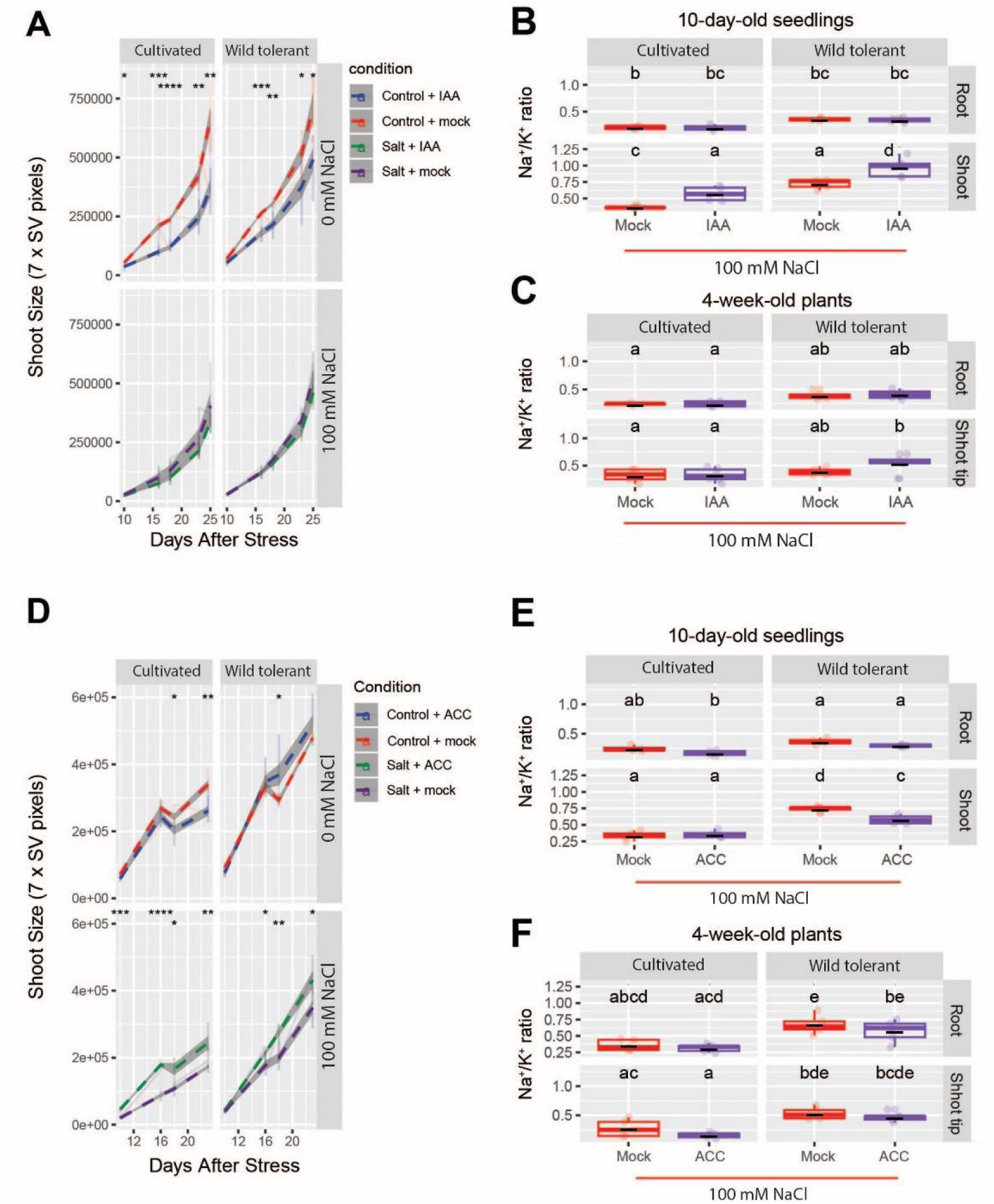
Foliar application of IAA and ACC increases and decreases the shoot Na⁺/K⁺ ratio shortly after salt stress exposure. Shoot size was monitored for foliar application of IAA **(A)** and ACC **(D)** over the period of 16 days in soil. The measurements were done based on 7-side view image pixels collected 10, 16, 18, and 23 days after salt stress, as indicated in the figure. The seeds were germinated in 1/4 MS media in the plates for 4 complete days. At d5, the seedlings were transplanted into the soil with 50% water holding capacity that contained 0 or 100 mM NaCl for 4 weeks. We initiated the application of hormones via foliar spray for a continuous period of five days following the transplantation into saline soil, followed by twice-per-week application until the end of the experiment, which was 4 weeks. The plants were imaged using PhenoCage setup, starting from 10 days after salt stress. KOH solution without IAA and water were used as mock treatments for IAA and ACC foliar applications, respectively. The asterisks above the graph in (A) and (D) indicate significant differences between no hormone treatment (mock) and hormone treatment conditions, as determined by the Student‘s t-test: *P < 0.05, **P < 0.01, ***P < 0.001, and ****P < 0.0001. Na^+^/K^+^ ratio of roots and shoots of two accessions after 10 days of salt stress for IAA **(B)** and ACC **(E)** treatment. Na^+^/K^+^ ratio of roots and shoot tips of two accessions after 4 weeks of salt stress for IAA **(C)** and ACC **(F)** treatment. (B-C, E-F) Statistical analysis was done by comparison of the means for all pairs using Tukey–Kramer HSD test for Levels not connected by the same letter are significantly different (P < 0.05).

To assess the impact of foliar IAA application on Na⁺ accumulation during short- and long-term salt exposure, we measured Na⁺, K⁺, and the Na⁺/K⁺ ratio in plant tissues after 10 days and 4 weeks of salt stress. After 10 days, although Na⁺ and K⁺ levels varied, no significant changes were detected across treatment groups, accessions, or tissues (**Fig. S7B-D**). However, analysis of the Na⁺/K⁺ ratio revealed a significant increase in shoot Na⁺/K⁺ ratio in both accessions following IAA application compared to the mock treatment (KOH solution without IAA) (**Fig. 7B**). In contrast, after 4 weeks of salt stress, no significant differences in Na⁺, K⁺, or the Na⁺/K⁺ ratio were observed among the treatment groups and accessions (**Fig. 7C, Fig. S7E-G**). These findings suggest that IAA transiently disrupts ion homeostasis during early salt stress exposure, but its effects diminish over prolonged stress periods.

Foliar ACC application showed that ACC and ACC mock treatments (water) had no effect on shoot size in either accession under control conditions until 16 days after application. In cultivated tomato, ACC application significantly reduced shoot size at 18- and 24-days post-treatment compared to the ACC mock group (**Fig. 7D, Fig. S7H**). In contrast, ACC treatment increased shoot size in the wild tolerant accession under control conditions, with a significant increase observed at 18 days post-treatment (**Fig. 7D**). Under salt stress, ACC consistently promoted shoot growth in both accessions across nearly all time points compared to the ACC mock group (**Fig. 7D, Fig. S7H**), suggesting that ACC has a dynamic role in regulating growth, with effects that vary depending on the accession and environmental conditions.

To investigate the effects of foliar ACC application on Na⁺ accumulation during short- and long-term salt exposure, we measured Na⁺, K⁺, and the Na⁺/K⁺ ratio in plant tissues after 10 days and 4 weeks after salt stress exposure. After 10 days of ACC treatment, no significant changes were observed in Na⁺, K⁺, or the Na⁺/K⁺ ratio in roots and shoots of either accession compared to their respective control groups (**Fig. S7I-K**). The only notable change was a significant reduction in the Na⁺/K⁺ ratio in the shoots of the wild tolerant accession under salt stress compared to the ACC mock treatment (**Fig. 7E**). After 4 weeks of ACC treatment, no significant differences in Na⁺, K⁺, or the Na⁺/K⁺ ratio were observed in the roots and shoots of either accession (**Fig. 7F, Fig. S7L-N**), except in the shoot tips of cultivated tomato, where both ACC and ACC mock treatments maintained significantly lower Na⁺ levels compared to the salt-only treatment (S+noACC) (**Fig. S7L**). These results suggest that while ACC treatment has some localized effects on Na⁺ accumulation, its impact on ion homeostasis is minimal over extended salt stress exposure.

### Root-specific transcriptomics points to changes in various hormonal-related genes under salt stress

To investigate the gene regulatory networks that reshape RSA in tomato under salt stress, we previously conducted RNA-Seq on root samples of both cultivated and wild-tolerant tomatoes following salt stress exposure as described in (Rahmati Ishka *et al*., 2024). Given the pivotal role of hormonal genes in regulating various aspects of plant growth and development, including root architecture, we focused on hormone-related genes differentially expressed under salt stress in these two tomato accessions. We identified 44 hormonal-related genes whose expression was significantly altered under salt stress, either in both accessions or only one within at least one-time point (**Fig. S8**). The identified genes included eight ethylene-related signaling genes that belong to the ethylene-insensitive family protein, ethylene-forming enzyme, and ethylene-responsive element binding proteins. Although the expression patterns for these eight genes are similar (increased or decreased in both accessions), there is a difference between accessions in terms of time points when the most significant changes are observed.

Of those 44 hormonal-related genes, 26 belonged to the auxin signaling pathway, including SAUR-like auxin-responsive protein family (17 genes out of 26), four Auxin efflux carrier family proteins (four genes out of 26), Auxin-responsive GH3 family proteins (two genes), and Auxin-responsive family protein, auxin-regulated gene involved in organ size, and Dormancy/auxin associated family protein (one gene for each). Six genes showed significant changes related to gibberellin signaling upon salt stress exposure. These included three members of the gibberellin 2-oxidase, two members of the gibberellin 2-oxidase 8, and one gene belonging to the gibberellin-regulated family protein. We also have found a few changes in gene expression of cytokinin and ABA-related genes. These included cytokinin oxidase 3 and 5, cytokinin oxidase/dehydrogenase 6, and abscisic acid responsive elements-binding factor 2. Together, these results highlight the importance of hormonal signaling in the maintenance of RSA under salt stress.

### Arabidopsis orthologs of tomato auxin-related genes exhibit altered root architecture and Na⁺ accumulation under salt stress

Because both agar-plate and foliar IAA applications increased Na⁺ accumulation in tomato accessions, we chose to further explore its role in Na⁺ accumulation by investigating Arabidopsis orthologs to the tomato auxin-related genes identified in our RNA-Seq data, leveraging the available genetic resources in Arabidopsis (**Fig. S8**). We obtained two independent alleles for auxin-related genes: PIN-LIKES 5 (*PILS5)*, which encodes an auxin efflux carrier family protein, and SMALL AUXIN UPREGULATED RNA 45 (*SAUR45)*, which encodes a SAUR-like auxin-responsive protein family. We examined the RSA of these mutants under control and salt stress conditions and evaluated their Na^+^ and K^+^ content (**Fig. 8, Fig. S9**). Neither of the two auxin mutants exhibited differences in main root length compared to Col-0 under control conditions (**Fig. 8A**). Under salt stress, both *saur45* mutant alleles showed a significant reduction in main root length relative to Col-0 (**Fig. 8A**). One allele from each mutant, *pils5-1* and *saur45-1*, displayed a significant decrease in lateral root growth under control conditions, their lateral root length remained unchanged under salt stress compared to Col-0 (**Fig. 8B**). Lateral root number remained consistent across genotypes and conditions, except for the *saur45-1* mutant, which exhibited a significant reduction in lateral root numbers under salt stress compared to Col-0 (**Fig. 8C**). These results suggest that auxin transport via PILS5 and SAUR45 signaling is key to root growth under salt stress. SAUR45 affects both main and lateral roots, while PILS5 supports lateral root growth only under control conditions.

**Figure 8.**
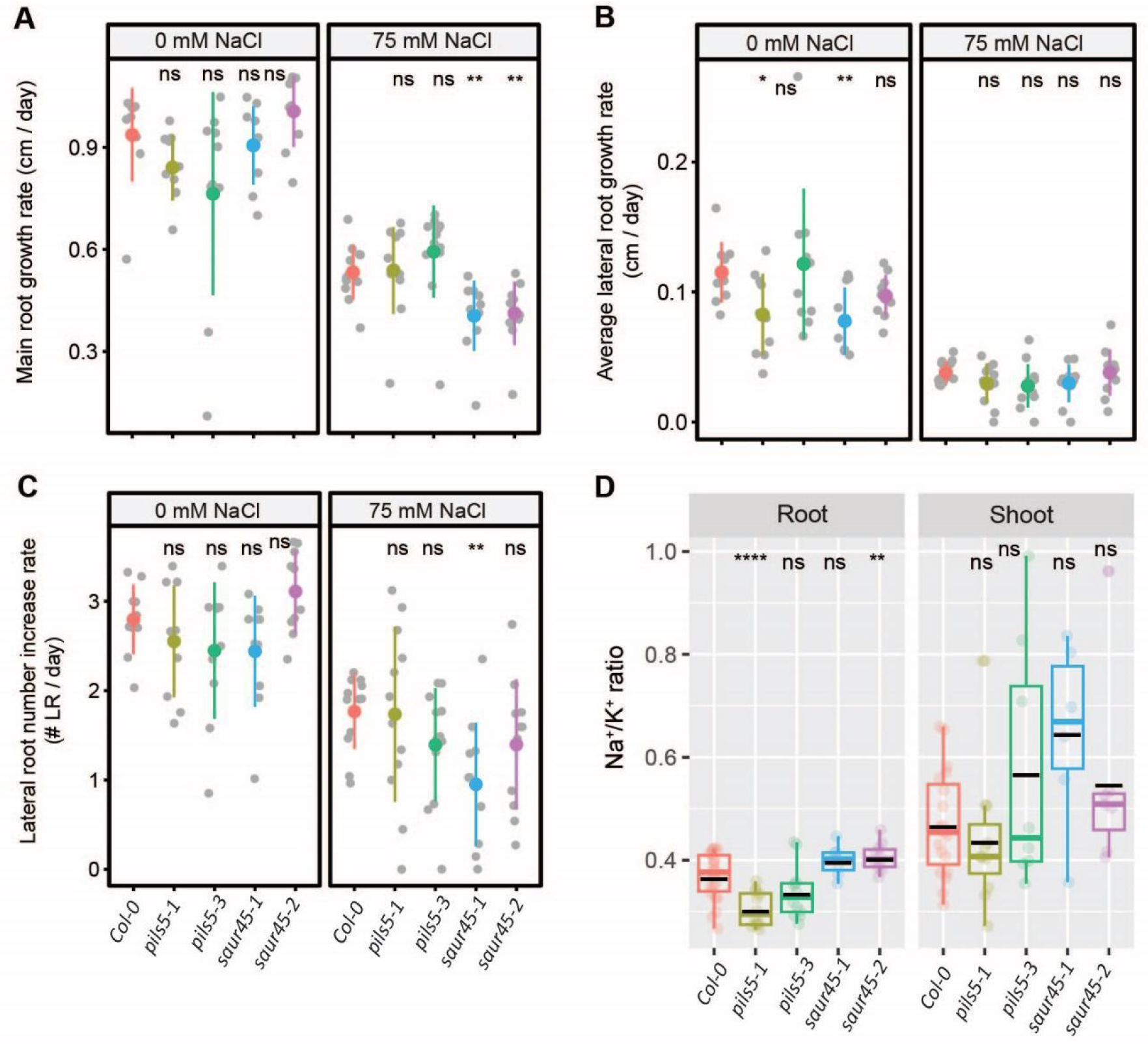
Arabidopsis *pils5* and *saur45* mutants, orthologs of tomato auxin-related genes, exhibit contrasting root Na⁺/K⁺ ratios. Root system architecture analysis of Arabidopsis Col-0 and auxin-related mutants under 0 or 75 mM concentrations of NaCl, as indicated in the figure, are shown here for main root growth rate **(A)**, average lateral root growth **(B)**, and number of lateral roots **(C)**. Na^+^/K^+^ ratio **(D)** of roots and shoots of different genotypes after 14 days on treatment plates. The asterisks above the graphs in (A-D) indicate significant differences between Col-0 wild-type and other genotypes by the Student‘s t-test: *P < 0.05, **P < 0.01, ***P < 0.001, and ****P < 0.0001, while "ns" indicates no significant difference.

To evaluate the impact of auxin-related gene loss on Na⁺ and K⁺ accumulation under salt stress, we measured Na⁺, K⁺, and the Na⁺/K⁺ ratio in roots and shoots after 10 days of salt treatment. Na⁺ levels were significantly reduced in both *pils5* mutant alleles in roots compared to Col-0, whereas no change was observed for *saur45* mutants (**Fig. S9A**). In shoots, Na⁺ levels were consistent across all genotypes except for *saur45-1*, which exhibited a significant increase compared to Col-0 (**Fig. S9A**). K⁺ levels remained stable across all genotypes and conditions (**Fig. S9B**). Notably, *pils5-1* and *saur45-2* displayed contrasting trends in the Na⁺/K⁺ ratio in roots, with a significant decrease and increase, respectively, while the remaining genotypes maintained a ratio similar to Col-0 in both roots and shoots (**Fig. 8D**). These results suggest that auxin signaling, particularly through PILS5 and SAUR45, plays a role in regulating Na⁺ homeostasis, with PILS5 influencing Na⁺ exclusion in roots, while SAUR45 triggers Na⁺ accumulation in shoots without altering root Na⁺ levels.

### Arabidopsis orthologs of tomato ethylene-related genes exhibit root growth sensitivity to salt stress

Since agar-plate and foliar ACC applications had differing effects on Na⁺ accumulation, showing either no change or a reduction in tomato accessions, respectively, we chose to investigate further ethylene’s role in Na⁺ accumulation by investigating Arabidopsis orthologs to the tomato ethylene-related genes identified in our RNA-Seq data (**Fig. S8**). We obtained two independent alleles for ethylene-related genes: *EIL4*, which encodes an ethylene-insensitive 3 family protein and *EGY2*, which encodes an ethylene-dependent gravitropism-deficient and yellow-green-like 2 protein. We examined the RSA of these mutants under control and salt stress conditions and evaluated their Na^+^ and K^+^ content (**Fig. 9, Fig. S10**). Our results indicated that under control conditions, none of the ACC-related mutants in Arabidopsis exhibited differences in main root growth compared to Col-0 (**Fig. 9A**). Interestingly, both alleles of the *egy2* mutant showed a significant reduction in main root length under salt stress compared to Col-0 (**Fig. 9A**). Additionally, both *eil4* and *egy2* mutants displayed a significant decrease in lateral root length under control conditions compared to Col-0, whereas only *eil4* mutants maintained this reduction under salt stress compared to Col-0 (**Fig. 9B**). While lateral root number remained unchanged in all mutants under control conditions compared to Col-0, *egy2-2* mutant showed a significant reduction (**Fig. 9C**). Under salt stress, only *eil4-2* and *egy2-2* mutants exhibited a significant decrease in lateral root number compared to Col-0 (**Fig. 9C**). These results suggest that ethylene signaling, particularly through EGY2 and EIL4, plays a critical role in regulating root growth under salt stress, with EGY2 being essential for main root growth adaptation and both EGY2 and EIL4 contributing to lateral root development under both control and salt stress conditions.

**Figure 9.**
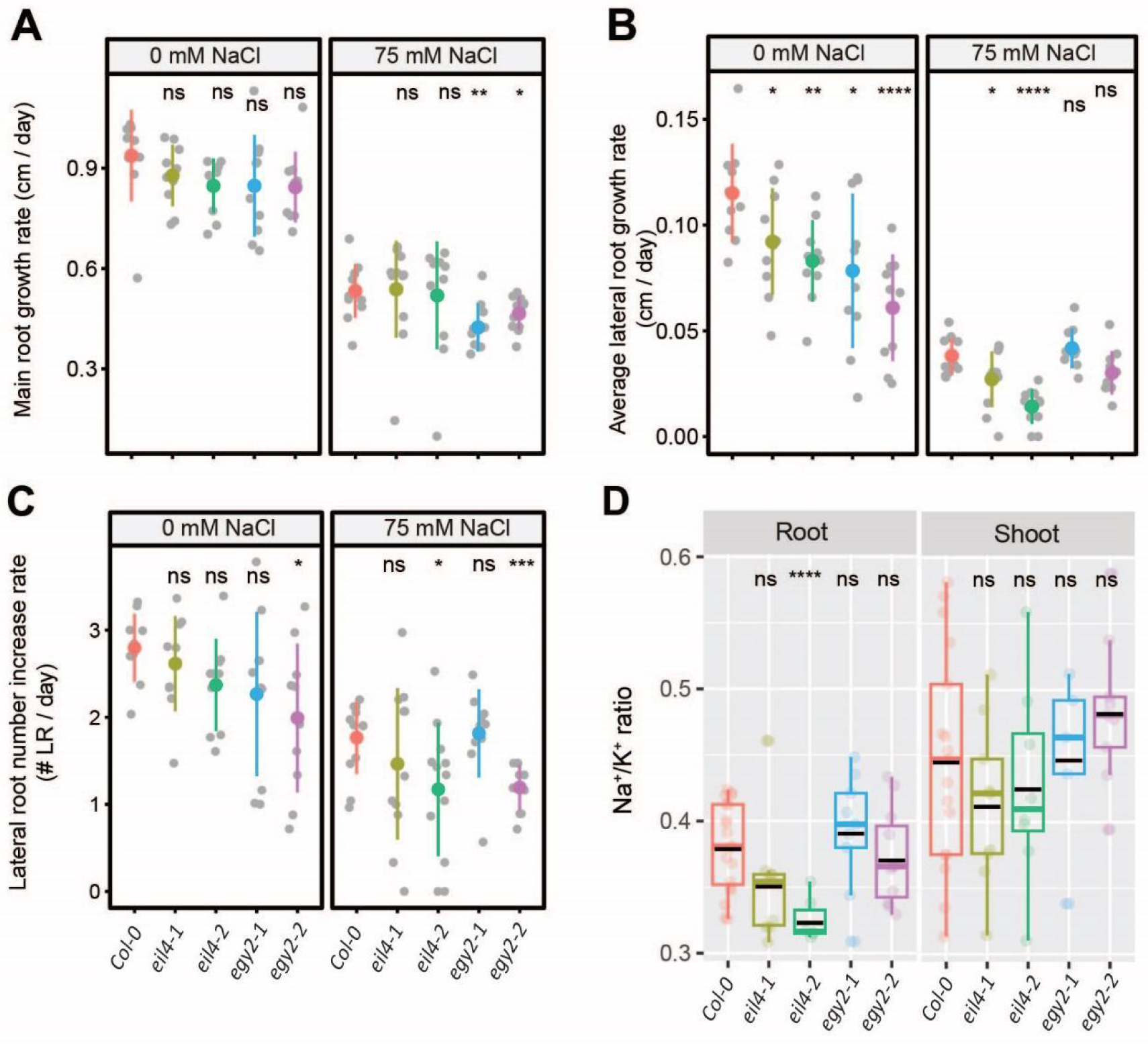
Arabidopsis *eil4*, but not *egy2*, orthologs of tomato ethylene-related genes, shows a reduced root Na⁺/K⁺ ratio. Root system architecture analysis of Arabidopsis Col-0 and ethylene-related mutants under 0 or 75 mM concentrations of NaCl, as indicated in the figure, are shown here for main root growth rate **(A)**, average lateral root growth **(B)**, and number of lateral roots **(C)**. Na^+^/K^+^ ratio **(D)** of roots and shoots of different genotypes after 14 days on treatment plates. The asterisks above the graphs in (A-D) indicate significant differences between Col-0 wild-type and other genotypes by the Student‘s t-test: *P < 0.05, **P < 0.01, ***P < 0.001, and ****P < 0.0001, while "ns" indicates no significant difference.

To evaluate the impact of ethylene-signaling-related gene loss on Na⁺ and K⁺ accumulation under salt stress, we measured Na⁺, K⁺, and the Na⁺/K⁺ ratio in roots and shoots after 10 days of salt treatment. Na⁺ levels remained largely unchanged across treatment groups and accessions, except in the roots of *eil4* mutants, which showed a significant reduction compared to Col-0 (**Fig. S10A**). Similarly, K⁺ levels remained unchanged across groups, except in the *eil4-1* mutant, which exhibited a significant decrease in root K⁺ levels compared to Col-0 (**Fig. S10B**). Among the tested mutants, only *eil4-2* showed a significantly reduced Na⁺/K⁺ ratio in roots compared to Col-0, while the other mutants showed ratios similar to Col-0 (**Fig. 9D**). These results suggest that ethylene signaling, specifically through EIL4, plays a role in regulating root Na⁺ exclusion, as disruption of EIL4 reduces Na⁺ accumulation and consequently lowers the Na⁺/K⁺ ratio in roots under salt stress.

### Defects in ethylene perception via tomato *nr* mutant promote main and lateral root elongation while reducing root Na⁺ accumulation and shoot K⁺ retention

Our analysis of tomato root architecture showed that ACC treatment inhibits several root traits but increases lateral root number. Unlike IAA, it does this without changing the Na⁺/K⁺ ratio. We examined the RSA of two available ethylene-related tomato mutants in control and salt stress conditions (**Fig. 10, Fig. S11**). One mutant, *never-ripe* (*nr*), carries a mutation in an ethylene receptor that causes dominant ethylene insensitivity, resulting in delayed fruit ripening and prolonged green fruit retention (Lanahan *et al*., 1994). The other mutant, *non-ripening ethylene insensitive* (*nei*), has a mutation in the *EIN2* ortholog, leading to recessive ethylene insensitivity (Guo *et al*., 2018). The *nr* mutant exhibited a significant increase in main root growth compared to its wild-type background, Ailsa Craig (AC), under both control and salt stress conditions, whereas *nei* showed no differences in main root length compared to its wild-type background, M82 (**Fig. 10A**). The *nr* mutant displayed a significant increase in lateral root length relative to AC under both control and salt stress conditions, whereas *nei* showed a significant reduction in lateral root length compared to M82 only under salt stress (**Fig. 10B**). The number of lateral roots remained unchanged across conditions, except for the *nr* mutant under salt stress, which exhibited a significant reduction compared to AC (**Fig. 10C**). Overall, the distinct root growth responses of the *nr* and *nei* mutants under salt stress suggest that ethylene perception through *nr* enhances main and lateral root elongation, whereas ethylene signaling via *nei* does not appear to play a significant role in regulating RSA under either non-stress or salt stress conditions.

**Figure 10.**
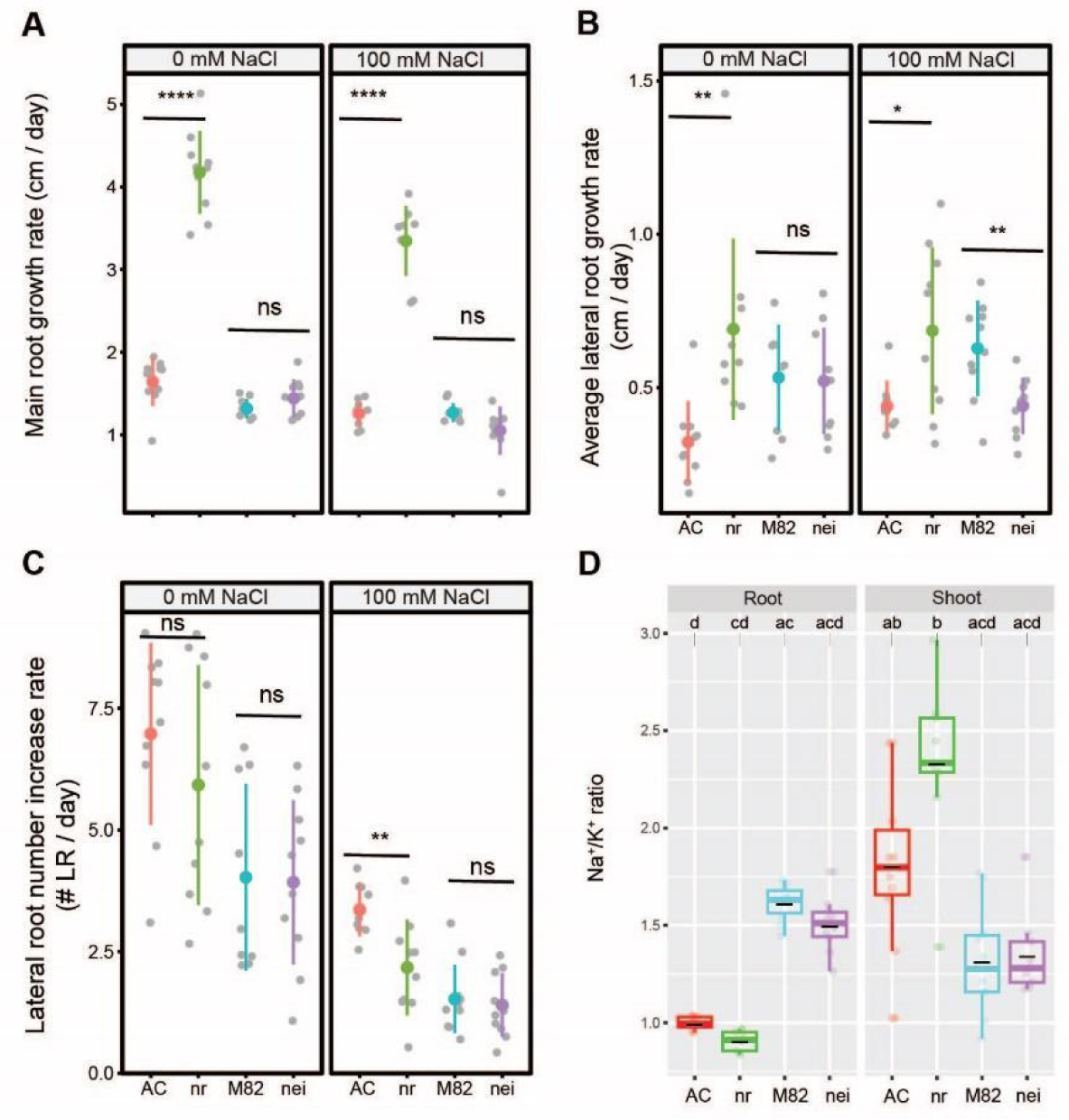
Tomato ethylene-related mutant *nr* exhibits increased root length in both main and lateral roots, along with an elevated shoot Na⁺/K⁺ ratio. Root system architecture analysis of tomato ethylene mutants along with their corresponding wild-type backgrounds under 0 or 100 mM concentrations of NaCl, as indicated in the figure, are shown here for main root growth rate **(A)**, average lateral root growth **(B)**, and number of lateral roots **(C)**. The asterisks above the graphs in (A-C) indicate significant differences between each mutant and its wild-type background by the Student‘s t-test: *P < 0.05, **P < 0.01, ***P < 0.001, and ****P < 0.0001, while "ns" indicates no significant difference. Na^+^/K^+^ ratio **(D)** of roots and shoots of different accessions after 10 days on treatment plates. Statistical analysis was done by comparison of the means for all pairs using Tukey–Kramer HSD test for (D). Levels not connected by the same letter are significantly different (P < 0.05). *nr* mutant is in the Ailsa Craig (AC) background and *nei* mutant is in the M82 background.

To evaluate the effect of tomato ethylene mutants on Na⁺ and K⁺ accumulation, we measured Na⁺, K⁺, and the Na⁺/K⁺ ratio in roots and shoots after 10 days of salt treatment (**Fig. 10D, Fig. S11A-B**). Both ethylene mutants accumulated similar levels of Na⁺ and K⁺ in roots and shoots compared to their respective wild-type backgrounds (**Fig. S11A-B**). The *nr* mutant exhibited a non-significant reduction in root Na^+^ and shoot K^+^ compared to its wild-type background. This resulted in an unchanged Na⁺/K⁺ ratio across all genotypes and tissues, although the *nr* mutant, maintained a non-significantly higher ratio in shoots compared to AC, mainly due to failure in K⁺ retention (**Fig. 10D**). These findings suggest that ethylene perception through *nr* can affect shoot ion homeostasis under salt stress, while ethylene signaling via *nei* does not appear to affect Na⁺ and K⁺ accumulation in roots or shoots.

## Discussion

Hormones play a key role in shaping root architecture by regulating meristem activity, lateral organ formation, and root growth (Ribba *et al*., 2020; Van Norman *et al*., 2013). This allows plants to adapt their roots to changing environments. Abscisic acid (ABA), auxins, cytokinins (CKs), gibberellic acid (GA), and ethylene are especially important under salt stress. Previous studies have explored how these hormones, alone (Fu *et al*., 2019; Qin *et al*., 2019; Li *et al*., 2014; Shan *et al*., 2014; Wang *et al*., 2015) or in combination (He *et al*., 2005; Moons *et al*., 1997; Lu *et al*., 2019), influence root development in saline conditions. While insightful, they vary widely in species, developmental stages, and experimental designs—making it hard to apply the findings broadly. Most focus only on primary root length (He *et al*., 2005), and few examine hormone effects on ion accumulation (Fu *et al*., 2019; Li *et al*., 2014). Here, we present a comprehensive study of how individual hormone treatments impact root architecture and ion accumulation under salt stress. We tested three tomato accessions—one cultivated and two wild—and uncovered distinct, species-specific hormonal responses.

Applying auxin, ACC, and gibberellin to tomato accessions showed that all three hormones boosted lateral root formation under salt stress (**Fig. 2, Fig. S2, Fig. 3, Fig. S3, Fig. 5, Fig. S5**). However, their effects on ion accumulation differed. Auxin increased shoot Na⁺/K⁺ ratios in wild tomatoes but not in the cultivated type (**Fig. 2**). ACC had no significant impact, while gibberellin lowered the ratio across all accessions (**Fig. 3, 5; Fig. S3, S5**). In Arabidopsis, auxin is known to suppress primary root growth while promoting lateral roots (Fu *et al*., 2019; He *et al*., 2005), which aligns with our observations. However, its impact on ion accumulation under salt stress has not been previously addressed in tomato. We found that auxin consistently increased lateral root numbers, regardless of salt treatment (**Fig. 2**). This may enhance Na⁺ uptake by increasing the number of entry points in the root system, as each lateral root must penetrate suberized barriers that are crucial for Na⁺ exclusion (Ursache *et al*., 2021; Tylová *et al*., 2017; B., Li *et al*., 2017) (Baxter *et al*., 2009). Notably, this effect was species-specific (**Fig. 2, Fig. S2**): wild tomatoes showed a higher Na⁺/K⁺ ratio, while the cultivated variety maintained a stable ratio—likely due to better K⁺ retention after auxin treatment.

In Arabidopsis, ACC application has been shown to inhibit primary root elongation (Qin *et al*., 2019), but its effects on other root traits and ion accumulation remained unexplored. In our study, ACC promoted lateral root formation in wild tomato accessions without altering Na⁺ or K⁺ levels (**Fig. 3, Fig. S3**). This was unexpected, as increased lateral root emergence was expected to be associated with enhanced Na⁺ uptake, as seen with auxin (**Fig. 2, Fig. S2**). These results suggest that the growth-promoting effects of ACC may be uncoupled from its effect on ion accumulation.

To evaluate our observations beyond agar plate-growth systems, we examined the long-term effects of auxin and ethylene on shoot growth and ion balance under salt stress. Foliar auxin application did not impact shoot size but increased the shoot Na⁺/K⁺ ratio during early salt exposure (day 10) in both accessions (**Fig. 7, Fig. S7**). In contrast, ACC (an ethylene precursor) promoted shoot growth and lowered the Na⁺/K⁺ ratio—but only in the wild-tolerant accession at the early stage of salt stress. This points to an accession-specific ethylene response. These findings suggest that ACC may affect ion balance more strongly in soil—where transpiration is higher than in agar-based systems.

Gibberellin treatment alone has been shown to increase primary root length and lateral root number in both *Arabidopsis* (Zhang *et al*., 2011) and rice (Qin *et al*., 2022), whereas it had the opposite effect in *Medicago*, where it reduced these root traits (Fonouni-Farde *et al*., 2019). Under salt stress, gibberellin had no effect on root length in dwarf OsGA2ox5-ox rice plants, which overexpressed the gibberellin 2-oxidase, a GA-deactivating gene (Shan *et al*., 2014). However, GA did restore the shoot height of GA2ox5-ox to wild-type levels (Shan *et al*., 2014). Our findings show that while GA₃ increased lateral root number, similar to auxin and ACC, it uniquely reduced the Na⁺/K⁺ ratio across all tomato accessions, primarily through enhancing potassium retention in both roots and shoots (**Fig. 5, Fig. S5**). This observation highlights the critical role of Gibberellin-mediated ion regulation as a key mechanism in improving salt stress resilience. The ability of GA₃ treatment to mitigate salt sensitivity in a wild-sensitive tomato accession suggests its potential as an exogenous intervention to alleviate salt stress across a broad spectrum of tomato varieties, including both cultivated and wild accessions with varying levels of salt tolerance.

Cytokinin treatment alone has been shown to suppress all aspects of root system development in both *Arabidopsis* (Beemster and Baskin, 2000) and maize (Márquez *et al*., 2019), as well as under salt stress in *Arabidopsis* (Wang *et al*., 2015). Similarly, abscisic acid (ABA) treatment alone reduces all root system components in *Arabidopsis* (Xing *et al*., 2016). Consistent with these findings, our study demonstrated that both ABA (**Fig. 4, Fig. S4**) and cytokinin (**Fig. 6, Fig. S6**) treatments negatively affected root growth traits across studied tomato accessions. Additionally, both hormones increased the Na⁺/K⁺ ratio in an accession-dependent manner. ABA elevated shoot Na⁺/K⁺ ratio in all accessions except the wild tolerant one (**Fig. 4**), while cytokinin increased it only in the cultivated tomato (**Fig. 6**). These results suggest that while ABA and cytokinin may have limited value in improving salt tolerance in cultivated tomatoes, they could offer potential benefits in stress mitigation for wild tomato-derived germplasm, such as newly domesticated *S. pimpinellifolium* lines (Zsögön *et al*., 2018).

Through functional characterization of Arabidopsis orthologs of tomato ethylene- and auxin-related genes (**Fig. 8, Fig. S9, Fig. 9, Fig. S10**), we further dissected several less-characterized components of hormone homeostasis underlying salt stress responses, including *EIL4*, *EGY2*, *PILS5*, and *SAUR45*. On the auxin side, *PILS5*, an ER-localized auxin transporter, has been implicated in regulating nuclear auxin signaling and ER stress responses (Waidmann *et al*., 2023), but its connection to salt stress and ion homeostasis has not been investigated until now. Our results demonstrated that *PILS5* contributes to Na⁺ exclusion in roots (**Fig. S9**), highlighting its role in maintaining ion homeostasis under salt stress. Additionally, while *SAUR45* has been linked to salt stress responses in the context of bacterial inoculation (Zhang *et al*., 2022), its direct role in root growth and ion homeostasis has not been explored. We found that mutation of *SAUR45* increases the sensitivity of the root system to salt and leads to elevated Na⁺ accumulation in shoots under salt stress (**Fig. 8, Fig. S9**), revealing a new layer of auxin-mediated regulation in stress adaptation. Together, these findings uncover previously unrecognized roles of auxin transport and signaling components in coordinating root architecture and ion homeostasis during salt stress.

While *EIN3* and *EIL1* are well-established as major transcription factors in ethylene signaling (Zhu *et al*., 2011), the role of *EIL4* in ethylene-mediated salt stress responses has remained unexplored. Similarly, *EGY2*, a thylakoid membrane-localized protease involved in chloroplast development, has not been previously linked to salt stress tolerance (Adamiec *et al*., 2018). Our findings demonstrated that ethylene homeostasis, particularly through Arabidopsis EGY2 and EIL4, plays a critical role in regulating root growth under salt stress, with EGY2 being essential for main root growth adaptation and both EGY2 and EIL4 contributing to lateral root development under both control and salt stress conditions (**Fig. 9, Fig. S10**). We further showed ethylene homeostasis, specifically through EIL4, plays a role in regulating root Na⁺ exclusion (**Fig. S10**), as disruption of EIL4 reduces Na⁺ accumulation and consequently lowers the Na⁺/K⁺ ratio in roots under salt stress (**Fig. 9**). These roles could involve mechanisms related to ROS homeostasis or retrograde signaling, similar to the function of *EGY3* in enhancing salt stress tolerance by promoting Cu/Zn-SOD2 stability and H₂O₂-mediated chloroplastic retrograde signaling (Zhuang *et al*., 2021). Unlike the *Arabidopsis* ethylene-related mutant *eil4*, the tomato ethylene-insensitive mutant *nr* exhibited enhanced root growth both in the presence and absence of salt (**Fig. 10**). However, it also showed increased Na⁺/K⁺ ratio in the shoot, mainly due to limiting K⁺ retention (**Fig. 10, Fig. S11**). This aligns with our observations from exogenous ACC application, which generally reduced Na⁺ accumulation in tomato. These results suggest that while ethylene homeostasis plays a conserved role in mediating salt resilience among tomato accessions, its function may not be conserved between more distantly related species, such as *Arabidopsis* and tomato.

Our analysis combining physiological and genetic approaches reveals the power of species-specific hormone treatments to improve plant performance under salt stress. Hormone balance is key to shaping root development, but its impact on salt tolerance varies by hormone, genotype, and species. Auxin, ACC, and gibberellin all promote lateral root growth, yet they affect ion accumulation in different ways. This points to hormone-specific roles in salt stress responses. Breaking salt stress responses into three parts—main root elongation, lateral root growth, and lateral root emergence—offers useful insight. However, root growth traits alone do not tell the whole story. To truly understand salt stress tolerance, we also need to assess ion accumulation, especially Na⁺ levels. We can only evaluate the long-term impact of strategies aimed at boosting salt tolerance by linking root traits to ion balance. Our findings show the importance of linking developmental and physiological traits when using hormones to boost stress resilience. Hormone-based treatments face fewer regulatory hurdles than many new agronomic compounds. Many hormones and their synthetic analogs are already approved for agricultural use. This makes them a quick and practical solution for enhancing crop performance in saline soils. Approaches like these could play a crucial role in ensuring food security as the global population continues to rise.

## Author contributions

M.R.I. conceived the project, performed the experiments, analyzed data, and wrote the manuscript with contributions from all coauthors. E.C. and M.P. assisted with ICP-AES, M.M.J. conceived the project, providing infrastructure, funding and supervision.

## Acknowledgements

The authors would like to thank Trent Anthony Donaldson, Stacey Na, Yalmarie Numan-Vazquez for their assistance with root tracing. The authors would like to thank Professor James J. Giovannoni for providing us with tomato ethylene mutants. The authors would also like to thank the BTI greenhouse and support staff for their help. Funding for this project was provided through NSF Mathematical Biology #2244735 (MMJ). The majority of funding for this work was generously provided by BTI’s startup funds awarded to MMJ.

## Supplemental figures and tables

**Figure S1. Salt stress reduces shoot size and fresh weight while increasing evapotranspiration, ion leakage, and shoot Na**⁺ **accumulation in all three tomato accessions.** Na⁺ **(A)** and K⁺ **(B)** in root and shoot of different accessions after 10 days on treatment plates. **(C)** Shoot size was monitored over eight days in soil using seven-side view image pixels collected every other day, as shown in the figure. Seeds were germinated on ¼ MS plates for four days. On day 5, seedlings were transferred to ¼ MS plates containing 0 or 100 mM NaCl for one week. Afterward, they were transplanted into soil at 50% water holding capacity (WHC) with either 0 or 100 mM NaCl for 35 days. Imaging was performed using the PhenoCage setup (doi: 10.1093/plphys/kiae237) every other day from 28 to 35 days after salt exposure. **(C)** Shoot fresh weight was measured at the end of the experiment in four-week-old plants. **(D)** Evapotranspiration was estimated by daily pot weight measurements, adjusted to the reference weight of 50% WHC with either water or 100 mM NaCl solution, and calculated as the difference in weight between consecutive days. **(F)** Shoot ion leakage was assessed in four-week-old plants following one week of salt stress in plates and three weeks in soil. **(G)** Na⁺ and **(H)** K⁺ contents in three distinct shoot tissues of four-week-old plants after two weeks of salt stress in soil. (A-B, D-H) Statistical analysis was done by comparison of the means for all pairs using Tukey–Kramer HSD test for Levels not connected by the same letter are significantly different (P < 0.05). (C) Significant differences between control and salt-stressed plants were determined using a Student’s t-test, with **, ***, and **** indicating p-values of <0.01, <0.001, and <0.0001, respectively.

**Figure S2. IAA treatment raises the Na^+^ accumulation in shoots of wild tomatoes but not cultivated tomato.** Na⁺ **(A)** and K⁺ **(B)** contents of root and shoot of different accessions after 10 days on treatment plates. (A-B) Statistical analysis was done by comparison of the means for all pairs using Tukey–Kramer HSD test for Levels not connected by the same letter are significantly different (P < 0.05).

**Figure S3. ACC treatment causes a non-significant decrease in Na^+^ contents in the roots and shoots of tolerant accessions.** Na^+^ **(A)** and K^+^ **(B)** content of root and shoot of different accessions after 10 days on treatment plates. (A-B) Statistical analysis was done by comparison of the means for all pairs using Tukey–Kramer HSD test for Levels not connected by the same letter are significantly different (P < 0.05).

**Figure S4. ABA treatment causes a significant decrease in shoot Na^+^ contents of wild tolerant accession.** Na^+^ **(A)** and K^+^ **(B)** content of root and shoot of different accessions after 10 days on treatment plates. (A-B) Statistical analysis was done by comparison of the means for all pairs using Tukey–Kramer HSD test for Levels not connected by the same letter are significantly different (P < 0.05).

**Figure S5. GA3 treatment decreases Na^+^ content while increasing K^+^ retention.** Na^+^ **(A)** and K^+^ **(B)** content of root and shoot of different accessions after 10 days on treatment plates. (A-B) Statistical analysis was done by comparison of the means for all pairs using Tukey–Kramer HSD test for Levels not connected by the same letter are significantly different (P < 0.05).

**Figure S6. Cytokinin treatment increases shoot Na^+^ content in cultivated and wild-sensitive accessions but not in wild tolerant tomato.** Na^+^ **(A)** and K^+^ **(B)** content of root and shoot of different accessions after 10 days on treatment plates. (A-B) Statistical analysis was done by comparison of the means for all pairs using Tukey–Kramer HSD test for Levels not connected by the same letter are significantly different (P < 0.05).

**Figure S7. Foliar application of IAA in soil-grown plants shows no significant effect on shoot size, whereas ACC application promotes shoot growth under salt stress.** Shoot size was monitored for foliar application of IAA **(A)** and ACC **(H)** over a period of 16 days in soil. The measurements were done based on 7-side view image pixels collected 10, 16, 18, and 23 days after salt stress, as indicated in the figure. The seeds were germinated in 1/4 MS media in the plates for 4 complete days. At d5, the seedlings were transplanted into the soil with 50% water holding capacity that contained 0 or 100 mM NaCl for 4 weeks. We initiated the application of hormones via foliar spray for a continuous period of five days following the transplantation into saline soil, followed by twice-per-week application until the end of the experiment, which was 4 weeks. The plants were imaged using PhenoCage setup, starting from 10 days after salt stress. KOH solution without IAA and water were used as mock treatments for IAA and ACC foliar applications, respectively. The asterisks above the graph in (A) and (H) indicate significant differences between no hormone treatment (mock) and hormone treatment conditions, as determined by the Student‘s t-test: *P < 0.05, **P < 0.01, ***P < 0.001, and ****P < 0.0001. Na⁺ and K⁺ content, along with the Na⁺/K⁺ ratio in roots and shoots of two accessions after 10 days of salt stress for IAA **(B-D)** and ACC **(I-K)** treatments. Na⁺ and K⁺ content, along with the Na⁺/K⁺ ratio in roots and shoot tips of two accessions after 4 weeks of salt stress for IAA **(E-G)** and ACC **(L-N)** treatments. (B-G, I-N) Statistical analysis was done by comparison of the means for all pairs using Tukey–Kramer HSD test for Levels not connected by the same letter are significantly different (P < 0.05). c-iaa represents control with IAA, c-mock represents control with mock, and c-noiaa represents control without treatment. Similarly, s-iaa represents salt with IAA, s-mock represents salt with mock, and s-noiaa represents salt without treatment. c-acc represents control with ACC, c-mock represents control with mock, and c-noacc represents control without treatment. Similarly, s-acc represents salt with ACC, s-mock represents salt with mock, and s-noacc represents salt without treatment.

**Figure S8. Tomato root-specific transcriptomic analysis reveals alterations in various hormone signaling-related genes under salt stress.** The heatmap displays the average normalized expression of 44 hormone-related genes from root-specific transcriptomic analysis under salt stress in cultivated and wild-tolerant tomatoes across various time points following salt stress exposure (Rahmati Ishka et al. 2025), as indicated in the figure. The x-axis numbers represent the minutes of salt stress exposure. Red and blue in the heatmap indicate upregulation and downregulation, respectively. *S. lyc* and *S. pimp* denote cultivated and wild-tolerant tomatoes, respectively.

**Figure S9. Arabidopsis *pils5* and *saur45* mutants exhibit contrasting Na**⁺ **accumulation.** Na⁺ **(A)** and K⁺ **(B)** contents of roots and shoots of different genotypes after 14 days on treatment plates. The asterisks above the graphs in (A-B) indicate significant differences between Col-0 and other genotypes, as determined by the Student’s t-test: *P < 0.05, **P < 0.01, and ***P < 0.001, while "ns" indicates no significant difference.

**Figure S10. Arabidopsis *eil4* mutants exhibit reduced root Na**⁺ **accumulation.** Na⁺ **(A)** and K⁺ **(B)** contents of roots and shoots of different genotypes after 14 days on treatment plates. The asterisks above the graphs in (A-B) indicate significant differences between Col-0 and other genotypes, as determined by the Student’s t-test: *P < 0.05, **P < 0.01, and ***P < 0.001, while "ns" indicates no significant difference.

**Figure S11. The tomato *nr* mutant shows impaired shoot K⁺ retention under salt stress.** Na⁺ (**A**) and K⁺ **(B)** contents of roots and shoots of different accessions after 10 days on treatment plates. Statistical analysis was done by comparison of the means for all pairs using Tukey–Kramer HSD test for (A-B). Levels not connected by the same letter are significantly different (P < 0.05). *nr* mutant is in the Ailsa Craig (AC) background and *nei* mutant is in the M82 background.

**Table S1. List of oligos used for genotyping Arabidopsis mutants.**

## References

Adamiec, M., Misztal, L., Kosicka, E., Paluch-Lubawa, E. and Luciński, R. (2018) Arabidopsis thaliana egy2 mutants display altered expression level of genes encoding crucial photosystem II proteins. J. Plant Physiol., 231, 155–167. Available at: 10.1016/j.jplph.2018.09.010.

Baxter, I., Hosmani, P.S., Rus, A., et al. (2009) Root suberin forms an extracellular barrier that affects water relations and mineral nutrition in Arabidopsis. PLoS Genet., 5, e1000492. Available at: 10.1371/journal.pgen.1000492.

Beemster, G.T. and Baskin, T.I. (2000) Stunted plant 1 mediates effects of cytokinin, but not of auxin, on cell division and expansion in the root of Arabidopsis. Plant Physiol., 124, 1718–1727. Available at: 10.1104/pp.124.4.1718.

Bianco, C. and Defez, R. (2009) Medicago truncatula improves salt tolerance when nodulated by an indole-3-acetic acid-overproducing Sinorhizobium meliloti strain. J. Exp. Bot., 60, 3097–3107. Available at: 10.1093/jxb/erp140.

Bleecker, A.B. and Kende, H. (2000) Ethylene: a gaseous signal molecule in plants. Annu. Rev. Cell Dev. Biol., 16, 1–18. Available at: 10.1146/annurev.cellbio.16.1.1.

Casimiro, I., Beeckman, T., Graham, N., Bhalerao, R., Zhang, H., Casero, P., Sandberg, G. and Bennett, M.J. (2003) Dissecting Arabidopsis lateral root development. Trends Plant Sci., 8, 165–171. Available at: 10.1016/S1360-1385(03)00051-7.

Davies, P.J. (2010) Plant Hormones: Biosynthesis, Signal Transduction, Action! Springer.

Dinler, B. (2021) Simultaneous treatment of different gibberellic acid doses InducesIon accumulation and response mechanisms to salt damage in MaizeRoots. Journal of Plant Biochemistry & Physiology, 9, 1–11. Available at: 10.35248/2329-9029.21.9.258 [Accessed June 17, 2025].

Fonouni-Farde, C., Miassod, A., Laffont, C., Morin, H., Bendahmane, A., Diet, A. and Frugier, F. (2019) Gibberellins negatively regulate the development of Medicago truncatula root system. Sci. Rep., 9, 2335. Available at: 10.1038/s41598-019-38876-1.

Fu, Y., Yang, Y., Chen, S., Ning, N. and Hu, H. (2019) Arabidopsis IAR4 modulates primary root growth under salt stress through ROS-mediated modulation of auxin distribution. Front. Plant Sci., 10, 522. Available at: 10.3389/fpls.2019.00522.

Gehan, M.A., Fahlgren, N., Abbasi, A., et al. (2017) PlantCV v2: Image analysis software for high-throughput plant phenotyping. PeerJ, 5, e4088. Available at: 10.7717/peerj.4088.

Guo, X., Zhang, Y., Tu, Y., Wang, Y., Cheng, W. and Yang, Y. (2018) Overexpression of an EIN3-binding F-box protein2-like gene caused elongated fruit shape and delayed fruit development and ripening in tomato. Plant Sci., 272, 131–141. Available at: 10.1016/j.plantsci.2018.04.016.

He, X.-J., Mu, R.-L., Cao, W.-H., Zhang, Z.-G., Zhang, J.-S. and Chen, S.-Y. (2005) AtNAC2, a transcription factor downstream of ethylene and auxin signaling pathways, is involved in salt stress response and lateral root development: AtNAC2 transcription factor and its function. Plant J., 44, 903–916. Available at: 10.1111/j.1365-313X.2005.02575.x.

Ishka, M.R., Sussman, H., Hu, Y., et al. (2024) Natural variation in salt-induced changes in root:shoot ratio reveals SR3G as a negative regulator of root suberization and salt resilience in Arabidopsis. Available at: 10.7554/eLife.98896.1.

Julkowska, M.M., Hoefsloot, H.C.J., Mol, S., Feron, R., Boer, G.-J. de, Haring, M.A. and Testerink, C. (2014) Capturing Arabidopsis root architecture dynamics with ROOT-FIT reveals diversity in responses to salinity. Plant Physiol., 166, 1387–1402. Available at: 10.1104/pp.114.248963.

Julkowska, M.M., Koevoets, I.T., Mol, S., et al. (2017) Genetic Components of Root Architecture Remodeling in Response to Salt Stress. Plant Cell, 29, 3198–3213. Available at: 10.1105/tpc.16.00680.

Lanahan, M.B., Yen, H.C., Giovannoni, J.J. and Klee, H.J. (1994) The never ripe mutation blocks ethylene perception in tomato. Plant Cell, 6, 521–530. Available at: 10.1105/tpc.6.4.521.

Li, B., Kamiya, T., Kalmbach, L., et al. (2017) Role of LOTR1 in Nutrient Transport through Organization of Spatial Distribution of Root Endodermal Barriers. Curr. Biol., 27, 758–765. Available at: 10.1016/j.cub.2017.01.030.

Li, J., Jia, H. and Wang, J. (2014) cGMP and ethylene are involved in maintaining ion homeostasis under salt stress in Arabidopsis roots. Plant Cell Rep., 33, 447–459. Available at: 10.1007/s00299-013-1545-8.

Li, X., Chen, L., Forde, B.G. and Davies, W.J. (2017) The biphasic root growth response to abscisic acid in Arabidopsis involves interaction with ethylene and auxin signalling pathways. Front. Plant Sci., 8, 1493. Available at: 10.3389/fpls.2017.01493.

Lobet, G., Pagès, L. and Draye, X. (2011) A novel image-analysis toolbox enabling quantitative analysis of root system architecture. Plant Physiol., 157, 29–39. Available at: 10.1104/pp.111.179895.

Lu, C., Chen, M.-X., Liu, R., et al. (2019) Abscisic acid regulates auxin distribution to mediate maize lateral root development under salt stress. Front. Plant Sci., 10, 716. Available at: 10.3389/fpls.2019.00716.

Lv, S., Yu, D., Sun, Q. and Jiang, J. (2018) Activation of gibberellin 20-oxidase 2 undermines auxin-dependent root and root hair growth in NaCl-stressed Arabidopsis seedlings. Plant Growth Regul., 84, 225–236. Available at: 10.1007/s10725-017-0333-9.

Marhavý, P., Duclercq, J., Weller, B., et al. (2014) Cytokinin controls polarity of PIN1-dependent auxin transport during lateral root organogenesis. Curr. Biol., 24, 1031–1037. Available at: 10.1016/j.cub.2014.04.002.

Márquez, G., Alarcón, M.V. and Salguero, J. (2019) Cytokinin inhibits lateral root development at the earliest stages of lateral root primordium initiation in maize primary root. J. Plant Growth Regul., 38, 83–92. Available at: 10.1007/s00344-018-9811-1.

Moons, A., Prinsen, E., Bauw, G. and Van Montagu, M. (1997) Antagonistic effects of abscisic acid and jasmonates on salt stress-inducible transcripts in rice roots. Plant Cell, 9, 2243–2259. Available at: 10.1105/tpc.9.12.2243.

Muraro, D., Byrne, H., King, J. and Bennett, M. (2013) The role of auxin and cytokinin signalling in specifying the root architecture of Arabidopsis thaliana. J. Theor. Biol., 317, 71–86. Available at: 10.1016/j.jtbi.2012.08.032.

Papon, N. and Courdavault, V. (2022) ARResting cytokinin signaling for salt-stress tolerance. Plant Sci., 314, 111116. Available at: https://www.sciencedirect.com/science/article/pii/S0168945221003125?casa_token=hd-T1FEuwZIAAAAA:n5LPdqcGvsLg99YlUWLBPagEGb3CicVwD86d7REtwFvV8Oa86M6X9piS3BaIg_T8iQRtUaMe_Q#fig0005.

Qin, H., Pandey, B.K., Li, Y., et al. (2022) Orchestration of ethylene and gibberellin signals determines primary root elongation in rice. Plant Cell, 34, 1273–1288. Available at: 10.1093/plcell/koac008.

Qin, H., Wang, J., Chen, X., et al. (2019) Rice OsDOF15 contributes to ethylene-inhibited primary root elongation under salt stress. New Phytol., 223, 798–813. Available at: 10.1111/nph.15824.

Rahmati Ishka, M., Sussman, H., Zhao, J., et al. (2024) Dissecting the genetic regulation of lateral root development in tomato under salt stress. bioRxiv. Available at: 10.1101/2024.06.20.599848.

Ribba, T., Garrido-Vargas, F. and O’Brien, J.A. (2020) Auxin-mediated responses under salt stress: from developmental regulation to biotechnological applications. J. Exp. Bot., 71, 3843–3853. Available at: 10.1093/jxb/eraa241.

Rivas, M.Á., Friero, I., Alarcón, M.V. and Salguero, J. (2022) Auxin-Cytokinin Balance Shapes Maize Root Architecture by Controlling Primary Root Elongation and Lateral Root Development. Front. Plant Sci., 13, 836592. Available at: 10.3389/fpls.2022.836592.

Riyazuddin, R., Verma, R., Singh, K., Nisha, N., Keisham, M., Bhati, K.K., Kim, S.T. and Gupta, R. (2020) Ethylene: A master regulator of salinity stress tolerance in plants. Biomolecules, 10, 959. Available at: 10.3390/biom10060959.

Rowe, J.H., Topping, J.F., Liu, J. and Lindsey, K. (2016) Abscisic acid regulates root growth under osmotic stress conditions via an interacting hormonal network with cytokinin, ethylene and auxin. New Phytol., 211, 225–239. Available at: 10.1111/nph.13882.

Santner, A., Calderon-Villalobos, L.I.A. and Estelle, M. (2009) Plant hormones are versatile chemical regulators of plant growth. Nat. Chem. Biol., 5, 301–307. Available at: 10.1038/nchembio.165.

Shan, C., Mei, Z., Duan, J., Chen, H., Feng, H. and Cai, W. (2014) OsGA2ox5, a gibberellin metabolism enzyme, is involved in plant growth, the root gravity response and salt stress. PLoS One, 9, e87110. Available at: 10.1371/journal.pone.0087110.

Shkolnik-Inbar, D., Adler, G. and Bar-Zvi, D. (2013) ABI4 downregulates expression of the sodium transporter HKT1;1 in Arabidopsis roots and affects salt tolerance. Plant J., 73, 993–1005. Available at: 10.1111/tpj.12091.

Tavladoraki, P., Cona, A., Federico, R., Tempera, G., Viceconte, N., Saccoccio, S., Battaglia, V., Toninello, A. and Agostinelli, E. (2012) Polyamine catabolism: target for antiproliferative therapies in animals and stress tolerance strategies in plants. Amino Acids, 42, 411–426. Available at: 10.1007/s00726-011-1012-1.

Thole, J.M., Beisner, E.R., Liu, J., Venkova, S.V. and Strader, L.C. (2014) Abscisic acid regulates root elongation through the activities of auxin and ethylene in Arabidopsis thaliana. G3 (Bethesda), 4, 1259–1274. Available at: 10.1534/g3.114.011080.

Tylová, E., Pecková, E., Blascheová, Z. and Soukup, A. (2017) Casparian bands and suberin lamellae in exodermis of lateral roots: an important trait of roots system response to abiotic stress factors. Ann. Bot., 120, 71–85. Available at: 10.1093/aob/mcx047.

Ursache, R., De Jesus Vieira Teixeira, C., Dénervaud Tendon, V., et al. (2021) GDSL-domain proteins have key roles in suberin polymerization and degradation. Nat Plants, 7, 353–364. Available at: 10.1038/s41477-021-00862-9.

Van Norman, J.M., Xuan, W., Beeckman, T. and Benfey, P.N. (2013) To branch or not to branch: the role of pre-patterning in lateral root formation. Development, 140, 4301–4310. Available at: 10.1242/dev.090548.

Waidmann, S., Béziat, C., Ferreira Da Silva Santos, J., Feraru, E., Feraru, M.I., Sun, L., Noura, S., Boutté, Y. and Kleine-Vehn, J. (2023) Endoplasmic reticulum stress controls PIN-LIKES abundance and thereby growth adaptation. Proc. Natl. Acad. Sci. U. S. A., 120, e2218865120. Available at: 10.1073/pnas.2218865120.

Wang, X., Wen, H., Suprun, A. and Zhu, H. (2025) Ethylene signaling in regulating plant growth, development, and stress responses. Plants, 14. Available at: 10.3390/plants14030309.

Wang, Y., Shen, W., Chan, Z. and Wu, Y. (2015) Endogenous cytokinin overproduction modulates ROS homeostasis and decreases salt stress resistance in Arabidopsis Thaliana. Front. Plant Sci., 6, 1004. Available at: 10.3389/fpls.2015.01004.

Werner, T., Motyka, V., Laucou, V., Smets, R., Van Onckelen, H. and Schmülling, T. (2003) Cytokinin-deficient transgenic Arabidopsis plants show multiple developmental alterations indicating opposite functions of cytokinins in the regulation of shoot and root meristem activity. Plant Cell, 15, 2532–2550. Available at: 10.1105/tpc.014928.

Xing, L., Zhao, Y., Gao, J., Xiang, C. and Zhu, J.-K. (2016) The ABA receptor PYL9 together with PYL8 plays an important role in regulating lateral root growth. Sci. Rep., 6, 27177. Available at: 10.1038/srep27177.

Yang, C., Ma, B., He, S.-J., et al. (2015) MAOHUZI6/ETHYLENE INSENSITIVE3-LIKE1 and ETHYLENE INSENSITIVE3-LIKE2 regulate ethylene response of roots and coleoptiles and negatively affect salt tolerance in rice. Plant Physiol., 169, 148–165. Available at: 10.1104/pp.15.00353.

Yu, L. ’ang, Sussman, H., Khmelnitsky, O., Rahmati Ishka, M., Srinivasan, A., Nelson, A.D.L. and Julkowska, M.M. (2024) Development of a mobile, high-throughput, and low-cost image-based plant growth phenotyping system. Plant Physiol. Available at: 10.1093/plphys/kiae237.

Zhang, Y., Tian, Z., Xi, Y., Wang, X., Chen, S., He, M., Chen, Y. and Guo, Y. (2022) Improvement of salt tolerance of *Arabidopsis thaliana* seedlings inoculated with endophytic *Bacillus cereus* KP120. J. Plant Interact., 17, 884–893. Available at: 10.1080/17429145.2022.2111471.

Zhang, Z.-L., Ogawa, M., Fleet, C.M., et al. (2011) Scarecrow-like 3 promotes gibberellin signaling by antagonizing master growth repressor DELLA in Arabidopsis. Proc. Natl. Acad. Sci. U. S. A., 108, 2160–2165. Available at: 10.1073/pnas.1012232108.

Zhao, H., Ma, T., Wang, X., Deng, Y., Ma, H., Zhang, R. and Zhao, J. (2015) OsAUX1 controls lateral root initiation in rice (Oryza sativa L.): OsAUX1 controlling lateral root initiation. Plant Cell Environ., 38, 2208–2222. Available at: 10.1111/pce.12467.

Zhuang, Y., Wei, M., Ling, C., et al. (2021) EGY3 mediates chloroplastic ROS homeostasis and promotes retrograde signaling in response to salt stress in Arabidopsis. Cell Rep., 36, 109384. Available at: 10.1016/j.celrep.2021.109384.

Zhu, Z., An, F., Feng, Y., et al. (2011) Derepression of ethylene-stabilized transcription factors (EIN3/EIL1) mediates jasmonate and ethylene signaling synergy in Arabidopsis. Proc. Natl. Acad. Sci. U. S. A., 108, 12539–12544. Available at: 10.1073/pnas.1103959108.

Zsögön, A., Čermák, T., Naves, E.R., et al. (2018) De novo domestication of wild tomato using genome editing. Nat. Biotechnol., 36, 1211–1216. Available at: 10.1038/nbt.4272.

