## Supplemental Figures for "Exogenous Hormone Treatments Reveal Species-Specific Regulation of Individual Components of Root Architecture and Salt Ion Accumulation in Cultivated and Wild Tomatoes"

### Supplemental figures and tables

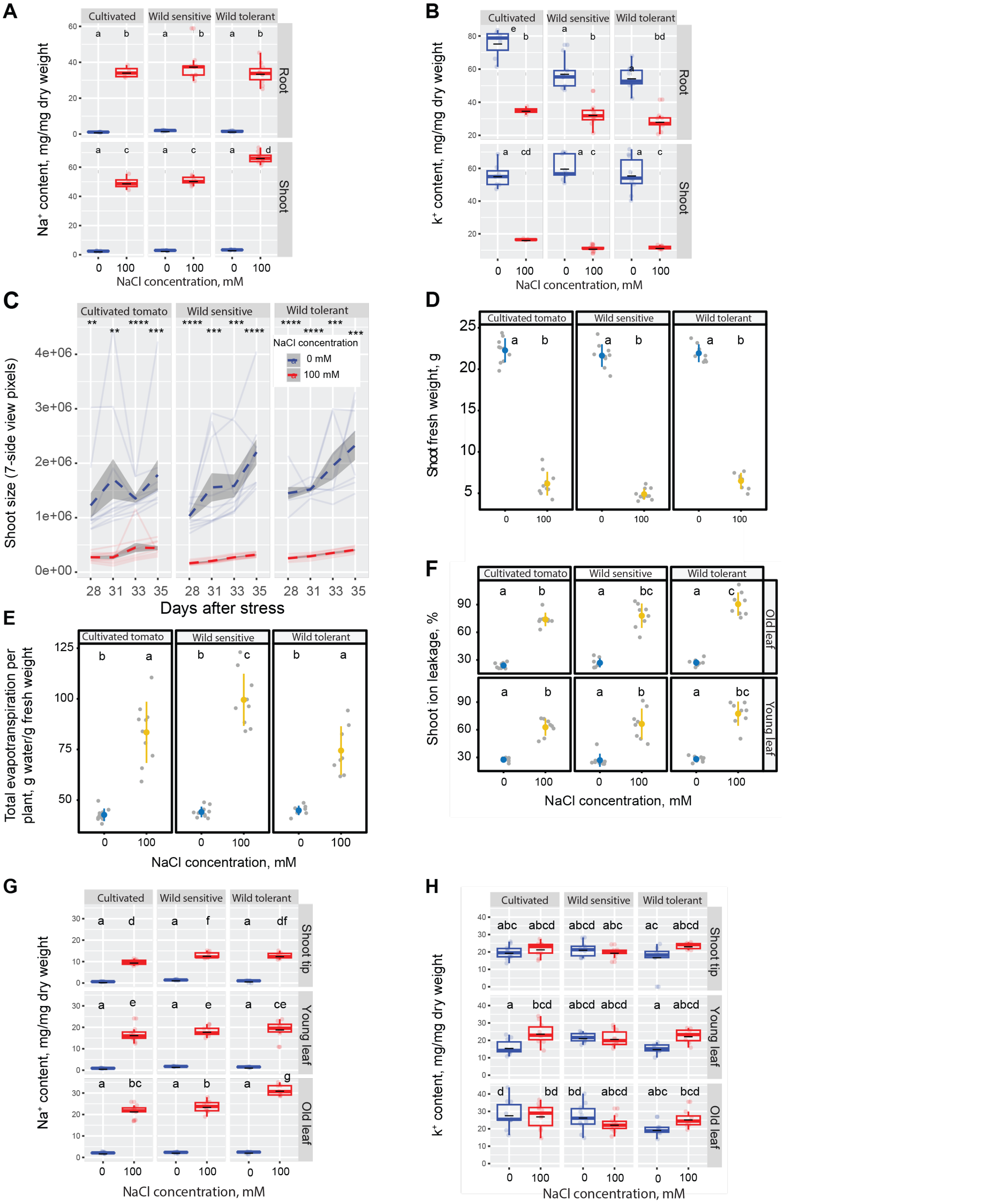


**Figure S1. Salt stress reduces shoot size and fresh weight while increasing evapotranspiration, ion leakage, and shoot $Na^+$ accumulation in all three tomato accessions.** $Na^+$ **(A)** and $K^+$ **(B)** in root and shoot of different accessions after 10 days on treatment plates. **(C)** Shoot size was monitored over eight days in soil using seven-side view image pixels collected every other day, as shown in the figure. Seeds were germinated on ¼ MS plates for four days. On day 5, seedlings were transferred to ¼ MS plates containing 0 or 100 mM NaCl for one week. Afterward, they were transplanted into soil at 50% water holding capacity (WHC) with either 0 or 100 mM NaCl for 35 days. Imaging was performed using the PhenoCage setup (doi: 10.1093/plphys/kiae237) every other day from 28 to 35 days after salt exposure. **(D)** Shoot fresh weight was measured at the end of the experiment in four-week-old plants. **(E)** Evapotranspiration was estimated by daily pot weight measurements, adjusted to the reference weight of 50% WHC with either water or 100 mM NaCl solution, and calculated as the difference in weight between consecutive days. **(F)** Shoot ion leakage was assessed in four-week-old plants following one week of salt stress in plates and three weeks in soil. **(G)** $Na^+$ and **(H)** $K^+$ contents in three distinct shoot tissues of four-week-old plants after two weeks of salt stress in soil. (A-B, D-H) Statistical analysis was done by comparison of the means for all pairs using Tukey–Kramer HSD test for Levels not connected by the same letter are significantly different ($P < 0.05$). (C) Significant differences between control and salt-stressed plants were determined using a Student's t-test, with **, ***, and **** indicating p-values of <0.01, <0.001, and <0.0001, respectively.

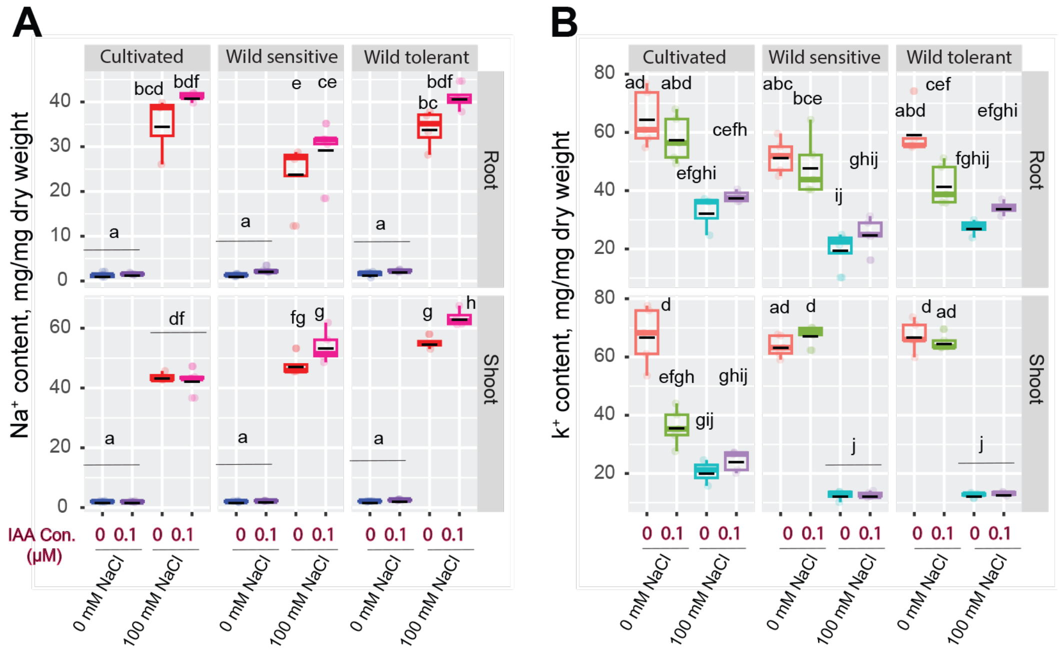


**Figure S2. IAA treatment raises the $Na^+$ accumulation in shoots of wild tomatoes but not cultivated tomato.** $Na^+$ **(A)** and $K^+$ **(B)** contents of root and shoot of different accessions after 10 days on treatment plates. (A-B) Statistical analysis was done by comparison of the means for all pairs using Tukey–Kramer HSD test for Levels not connected by the same letter are significantly different ($P < 0.05$).

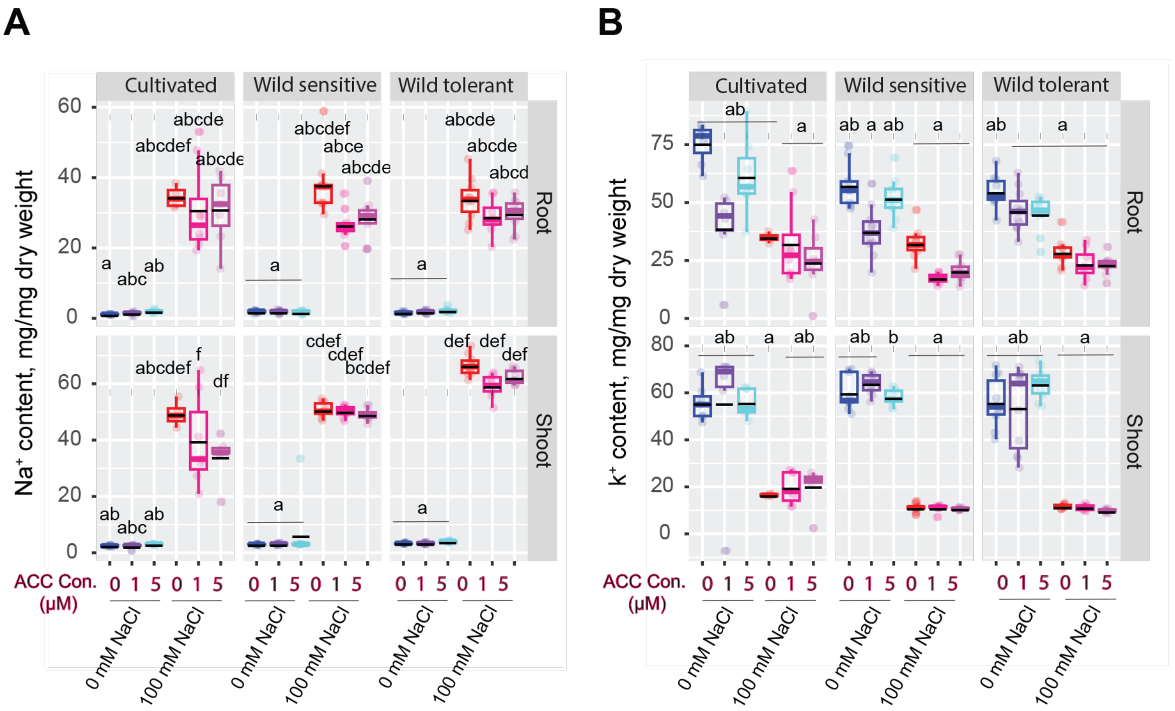


**Figure S3. ACC treatment causes a non-significant decrease in Na$^+$ contents in the roots and shoots of tolerant accessions.** Na$^+$ **(A)** and K$^+$ **(B)** content of root and shoot of different accessions after 10 days on treatment plates. (A-B) Statistical analysis was done by comparison of the means for all pairs using Tukey–Kramer HSD test for Levels not connected by the same letter are significantly different ($P < 0.05$).

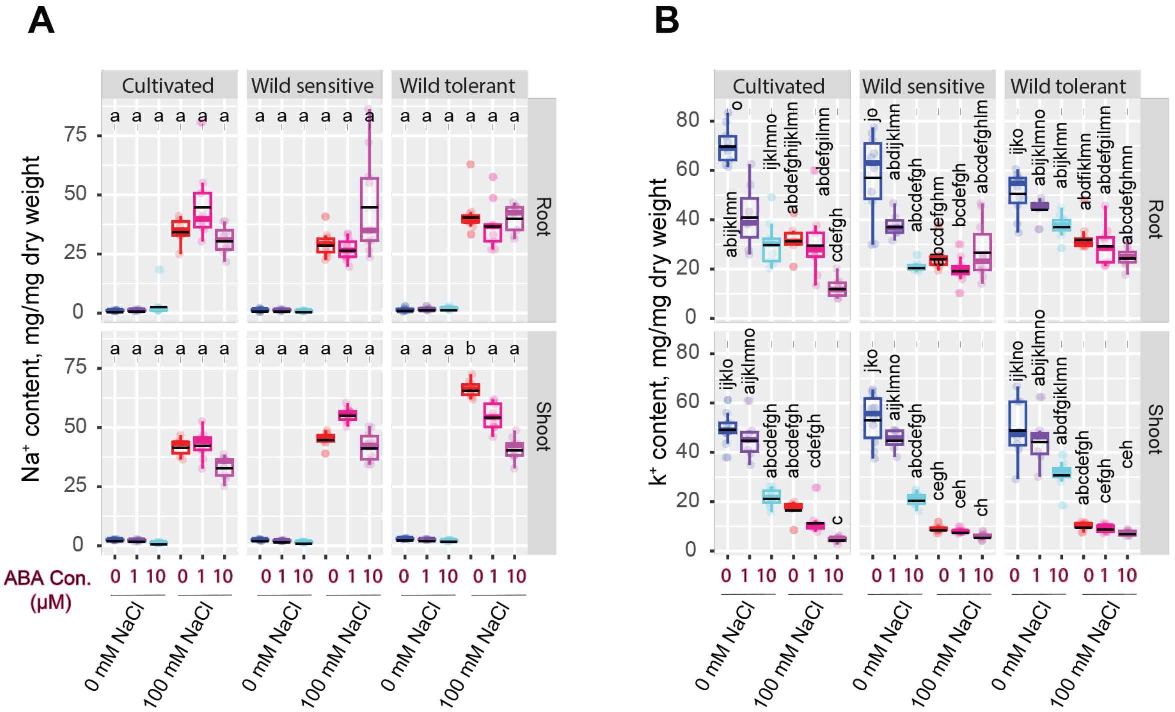


**Figure S4. ABA treatment causes a significant decrease in shoot Na$^+$ contents of wild tolerant accession.** Na$^+$ **(A)** and K$^+$ **(B)** content of root and shoot of different accessions after 10 days on treatment plates. (A-B) Statistical analysis was done by comparison of the means for all pairs using Tukey–Kramer HSD test for Levels not connected by the same letter are significantly different ($P < 0.05$).

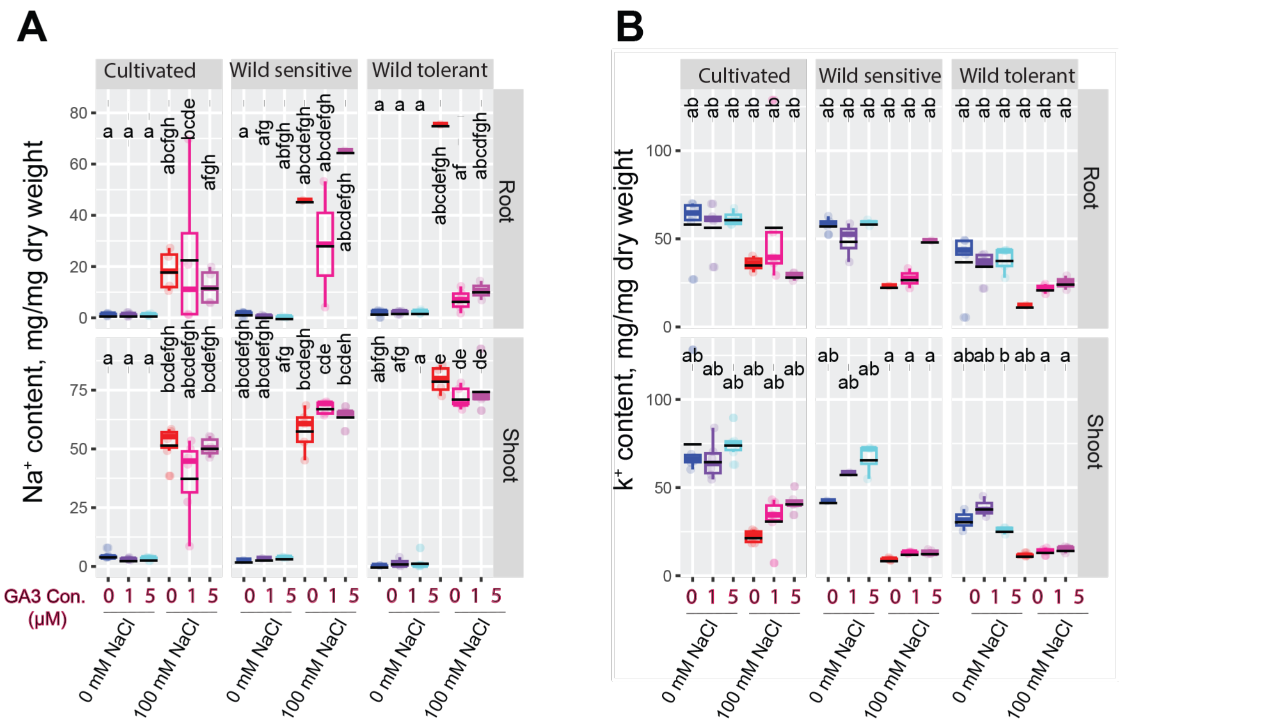


**Figure S5. GA3 treatment decreases $Na^+$ content while increasing $K^+$ retention.** $Na^+$ **(A)** and $K^+$ **(B)** content of root and shoot of different accessions after 10 days on treatment plates. (A-B) Statistical analysis was done by comparison of the means for all pairs using Tukey–Kramer HSD test for Levels not connected by the same letter are significantly different ($P < 0.05$).

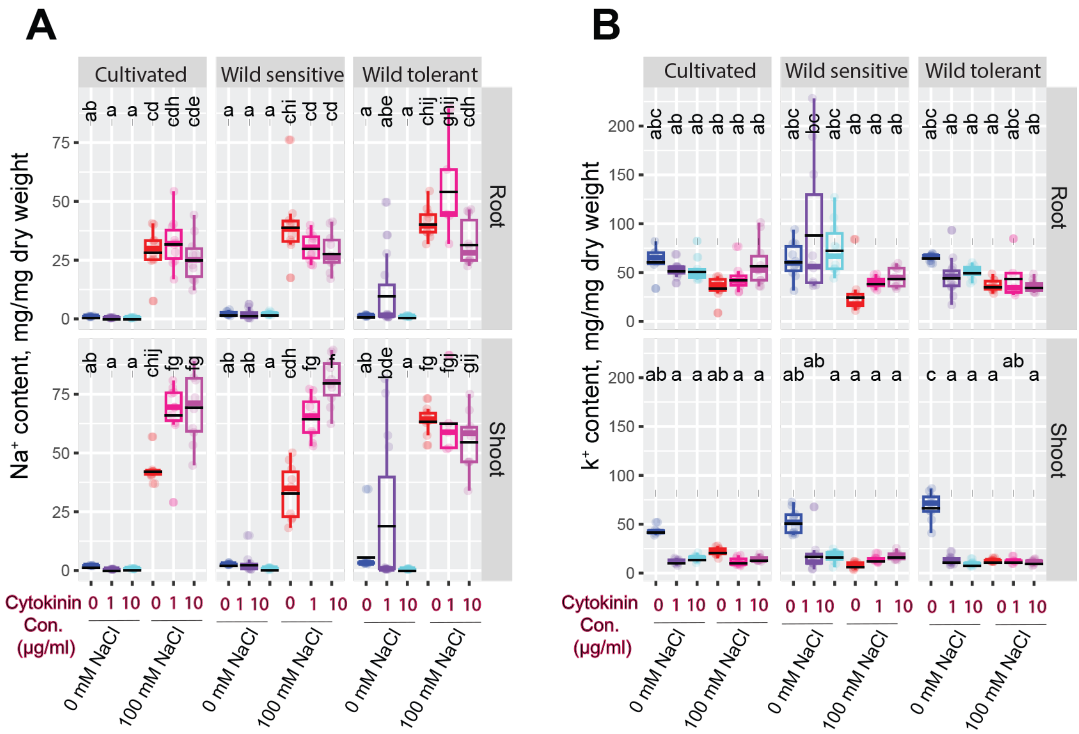


**Figure S6. Cytokinin treatment increases shoot $Na^+$ content in cultivated and wild-sensitive accessions but not in wild tolerant tomato.** $Na^+$ **(A)** and $K^+$ **(B)** content of root and shoot of different accessions after 10 days on treatment plates. (A-B) Statistical analysis was done by comparison of the means for all pairs using Tukey–Kramer HSD test for Levels not connected by the same letter are significantly different ($P < 0.05$).

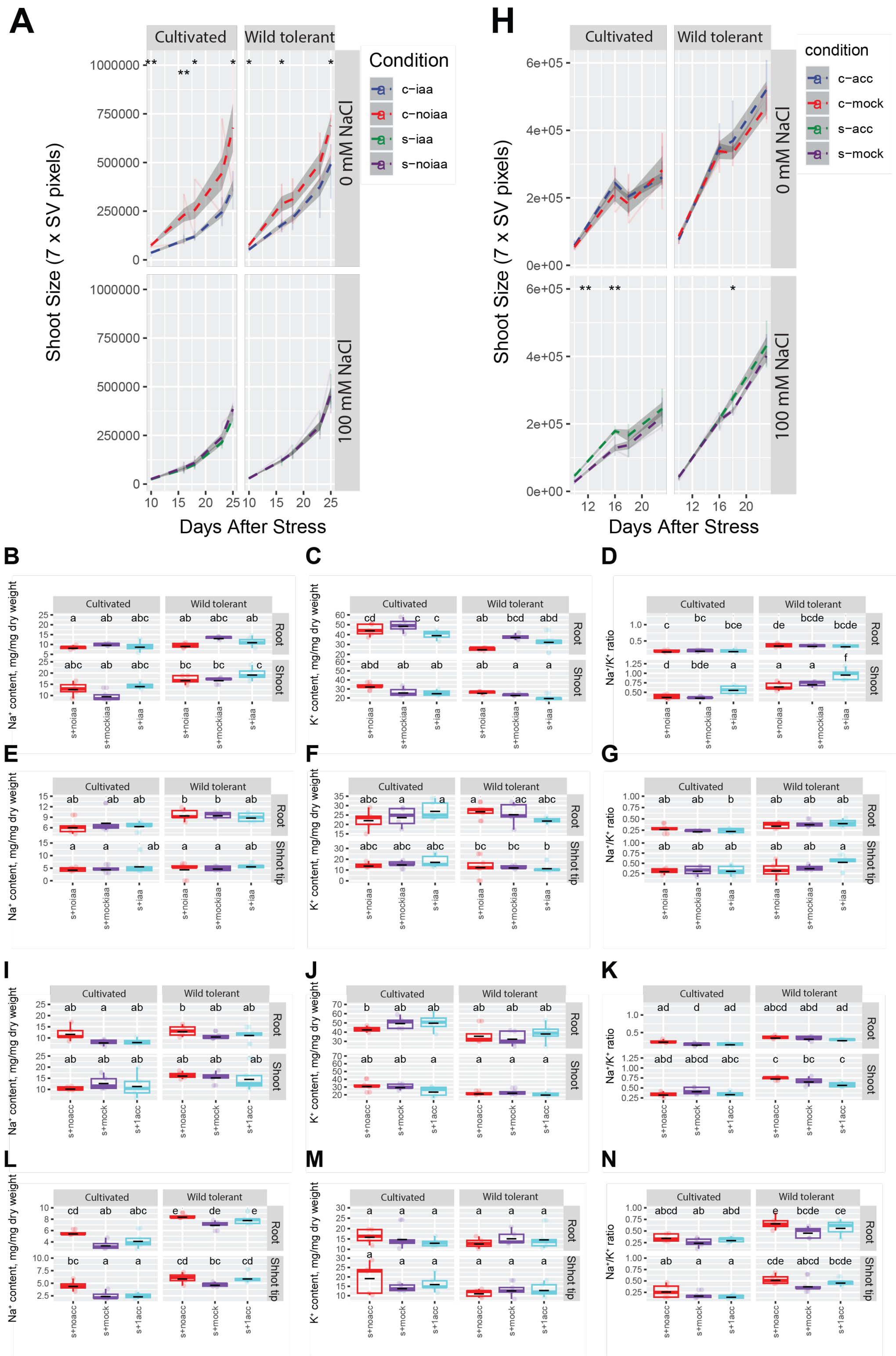


**Figure S7. Foliar application of IAA in soil-grown plants shows no significant effect on shoot size, whereas ACC application promotes shoot growth under salt stress.** Shoot size was monitored for foliar application of IAA **(A)** and ACC **(H)** over a period of 16 days in soil. The measurements were done based on 7-side view image pixels collected 10, 16, 18, and 23 days after salt stress, as indicated in the figure. The seeds were germinated in 1/4 MS media in the plates for 4 complete days. At d5, the seedlings were transplanted into the soil with 50% water holding capacity that contained 0 or 100 mM NaCl for 4 weeks. We initiated the application of hormones via foliar spray for a continuous period of five days following the transplantation into saline soil, followed by twice-per-week application until the end of the experiment, which was 4 weeks. The plants were imaged using PhenoCage setup, starting from 10 days after salt stress. KOH solution without IAA and water were used as mock treatments for IAA and ACC foliar applications, respectively. The asterisks above the graph in (A) and (H) indicate significant differences between no hormone treatment (mock) and hormone treatment conditions, as determined by the Student's t-test: *$P < 0.05$, **$P < 0.01$, ***$P < 0.001$, and ****$P < 0.0001$. $Na^+$ and $K^+$ content, along with the $Na^+/K^+$ ratio in roots and shoots of two accessions after 10 days of salt stress for IAA **(B-D)** and ACC **(I-K)** treatments. $Na^+$ and $K^+$ content, along with the $Na^+/K^+$ ratio in roots and shoot tips of two accessions after 4 weeks of salt stress for IAA **(E-G)** and ACC **(L-N)** treatments. (B-G, I-N) Statistical analysis was done by comparison of the means for all pairs using Tukey–Kramer HSD test for Levels not connected by the same letter are significantly different ($P < 0.05$). c-iaa represents control with IAA, c-mock represents control with mock, and c-noiaa represents control without treatment. Similarly, s-iaa represents salt with IAA, s-mock represents salt with mock, and s-noiaa represents salt without treatment. c-acc represents control with ACC, c-mock represents control with mock, and c-noacc represents control without treatment. Similarly, s-acc represents salt with ACC, s-mock represents salt with mock, and s-noacc represents salt without treatment.

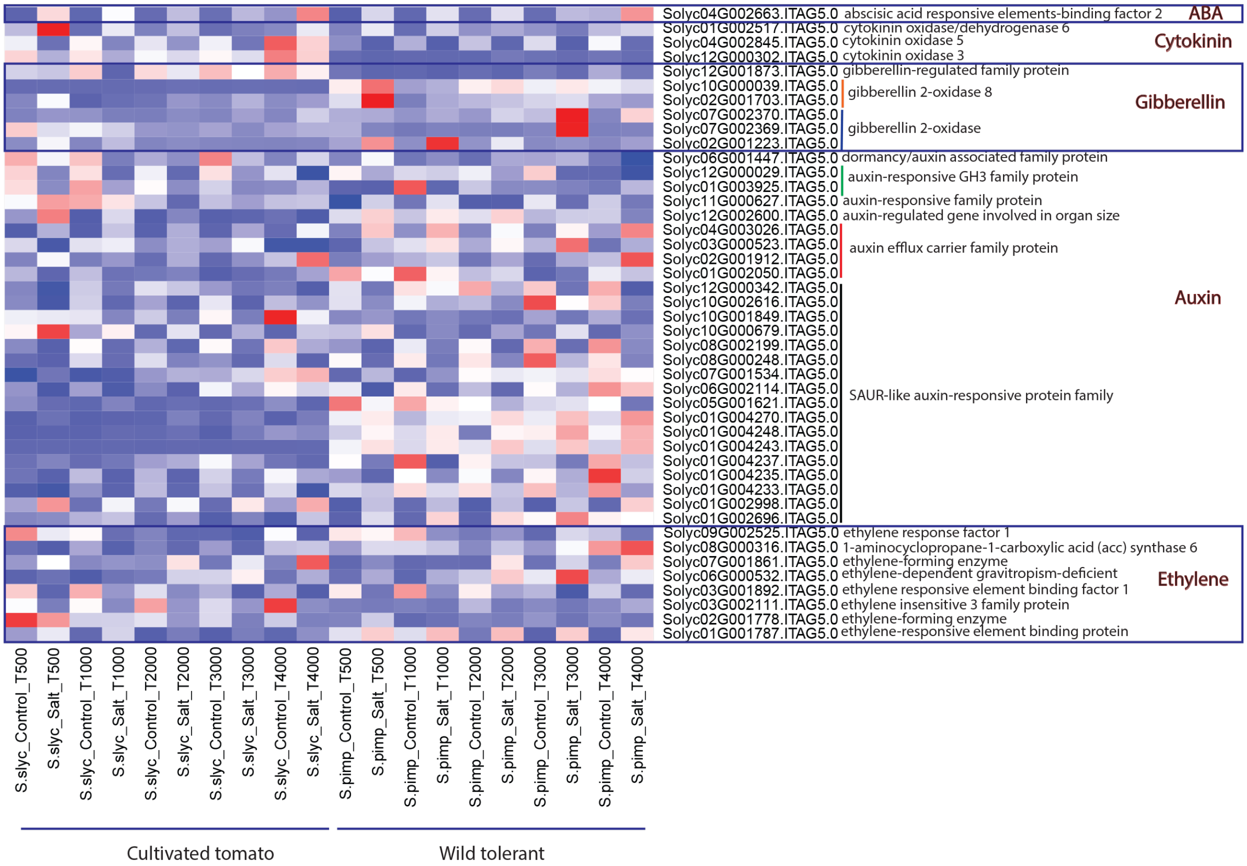


**Figure S8. Tomato root-specific transcriptomic analysis reveals alterations in various hormone signaling-related genes under salt stress.** The heatmap displays the average normalized expression of 44 hormone-related genes from root-specific transcriptomic analysis under salt stress in cultivated and wild-tolerant tomatoes across various time points following salt stress exposure (Rahmati Ishka et al. 2025), as indicated in the figure. The x-axis numbers represent the minutes of salt stress exposure. Red and blue in the heatmap indicate upregulation and downregulation, respectively. *S. lyc* and *S. pimp* denote cultivated and wild-tolerant tomatoes, respectively.

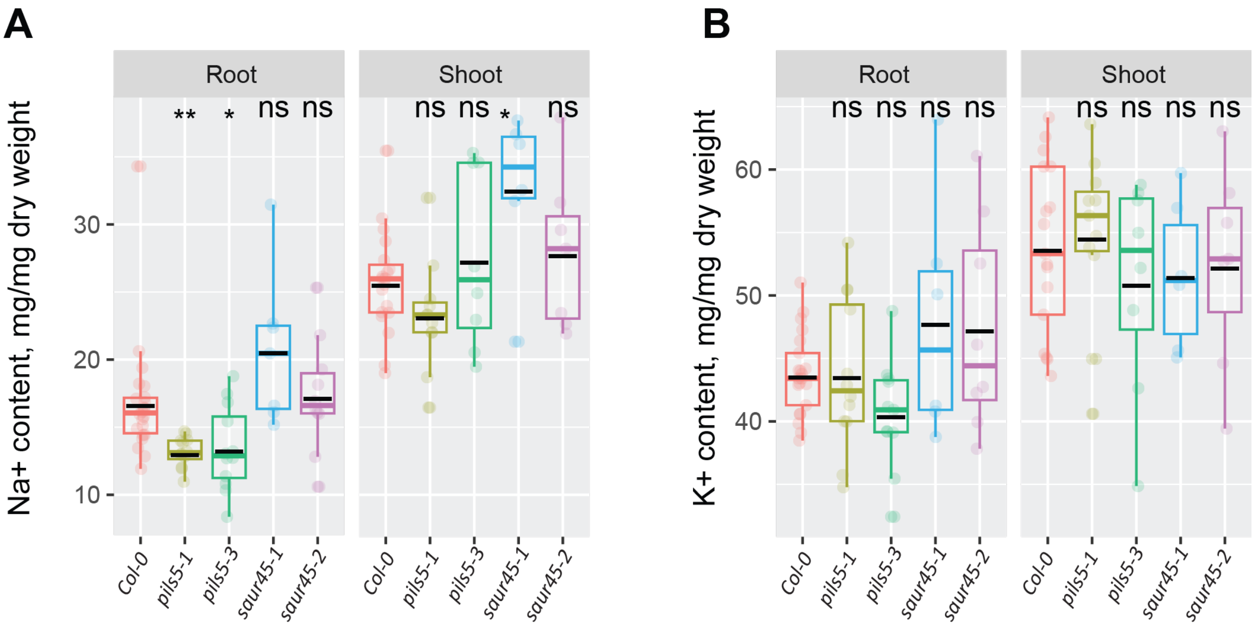


**Figure S9. Arabidopsis *pils5* and *saur45* mutants exhibit contrasting Na⁺ accumulation.** $Na^+$ **(A)** and $K^+$ **(B)** contents of roots and shoots of different genotypes after 14 days on treatment plates. The asterisks above the graphs in (A-B) indicate significant differences between Col-0 and other genotypes, as determined by the Student's t-test: *P < 0.05, **P < 0.01, and ***P < 0.001, while "ns" indicates no significant difference.

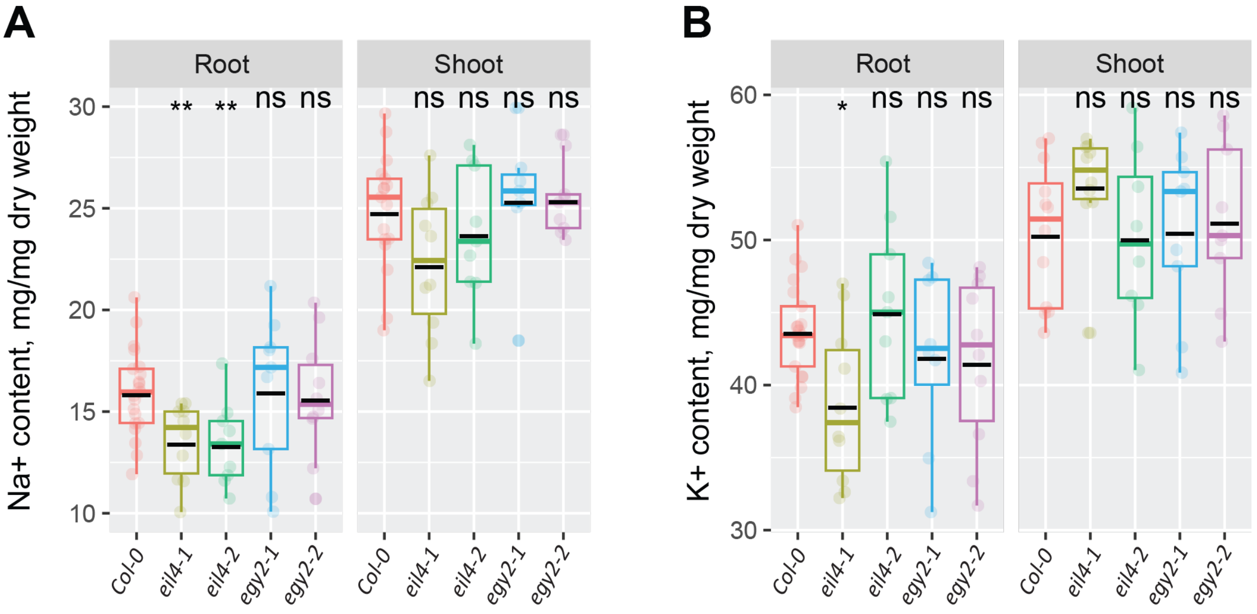


**Figure S10. Arabidopsis *eil4* mutants exhibit reduced root $Na^+$ accumulation.** $Na^+$ **(A)** and $K^+$ **(B)** contents of roots and shoots of different genotypes after 14 days on treatment plates. The asterisks above the graphs in (A-B) indicate significant differences between Col-0 and other genotypes, as determined by the Student's t-test: *P < 0.05, **P < 0.01, and ***P < 0.001, while "ns" indicates no significant difference.

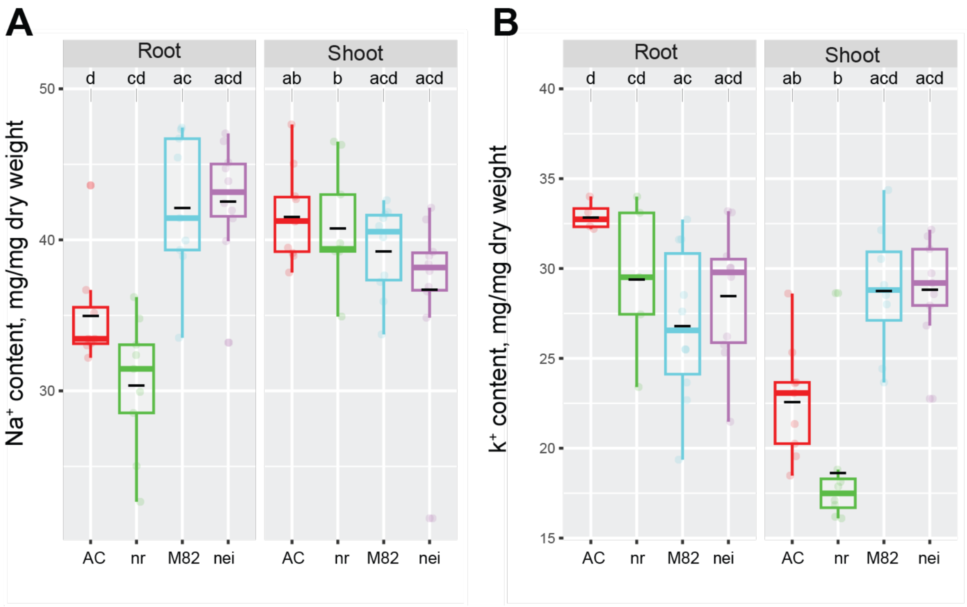


**Figure S11. The tomato *nr* mutant shows impaired shoot $K^+$ retention under salt stress.** $Na^+$ **(A)** and $K^+$ **(B)** contents of roots and shoots of different accessions after 10 days on treatment plates. Statistical analysis was done by comparison of the means for all pairs using Tukey–Kramer HSD test for (A-B). Levels not connected by the same letter are significantly different ($P < 0.05$). *nr* mutant is in the Ailsa Craig (AC) background and *nei* mutant is in the M82 background.

**Table S1. List of oligos used for genotyping Arabidopsis mutants.**
